# Mechanism of actin polymerization revealed by cryo-EM structures of actin filaments with three different bound nucleotides

**DOI:** 10.1101/309534

**Authors:** Steven Z. Chou, Thomas D. Pollard

## Abstract

We used electron cryo-micrographs to reconstruct actin filaments with bound AMPPNP (β,γ-imidoadenosine 5’-triphosphate, an ATP analog), ADP-P_i_ (ADP with inorganic phosphate) or ADP to resolutions of 3.4 Å, 3.4 Å and 3.6 Å. Subunits in the three filaments have nearly identical backbone conformations, so assembly rather than ATP hydrolysis or phosphate dissociation is responsible for their flattened conformation in filaments. Polymerization increases the rate of ATP hydrolysis by changing the conformations of the three ATP phosphates and the side chains of Gln137 and His161 in the active site. Flattening also promotes interactions along both the long-pitch and short-pitch helices. In particular, conformational changes in subdomain 3 open up favorable interactions with the DNase-I binding loop in subdomain 2 of the adjacent subunit. Subunits at the barbed end of the filament are likely to be in this favorable conformation, while monomers are not. This difference explains why filaments grow faster at the barbed end than the pointed end. Loss of hydrogen bonds after phosphate dissociation may account for the greater flexibility of ADP-actin filaments.

**Significance Statement:** Actin filaments comprise a major part of the cytoskeleton of eukaryotic cells and serve as tracks for myosin motor proteins. The filaments assemble from actin monomers with a bound ATP. After polymerization, actin rapidly hydrolyzes the bound ATP and slowly dissociates the γ-phosphate. ADP-actin filaments then disassemble to recycle the subunits. Understanding how actin filaments assemble, disassemble and interact with numerous regulatory proteins depends on knowing the structure of the filament. High quality structures of ADP-actin filaments were available, but not of filaments with bound ATP- or with ADP and phosphate. We determined structures of actin filaments with bound AMPPNP (a slowly hydrolyzed ATP analog), ADP and phosphate and ADP by cryo-electron microscopy. These structures show how conformational changes during actin assembly promote ATP hydrolysis and faster growth at one end of the filament than the other.

## Introduction

Actin, one of the most abundant proteins in eukaryotes, contributes to both cellular structure and motility. Filaments of actin form the cytoskeleton and interact with myosin motor proteins for cytokinesis, muscle contraction and transporting particles inside cells. Decades of biochemical analysis produced a detailed explanation actin assembly, including rate and equilibrium constants for most of the reactions (Dominguez and Holmes 2011, Pollard 2016). Actin binds ATP, which is hydrolyzed when the protein incorporates into filaments (Straub and Feuer 1950). Subsequent slow dissociation of the γ-phosphate (Carlier and Pantaloni 1986), favors depolymerization and changes the affinity of the filament for proteins such as cofilin (Pollard 2016).

In spite of more than 100 crystal structures of actin monomers (Fig. S1A) (Dominguez and Holmes 2011), the limited resolution of the available structures of actin filaments (von der Ecken, Müller et al. 2014, Galkin, Orlova et al. 2015, von der Ecken, Heissler et al. 2016) has left open many questions about all aspects of actin assembly. For example, why do filaments hydrolyze ATP thousands of times faster than monomers (Blanchoin and Pollard 2002, Rould, Wan et al. 2006), why do filaments elongate faster at their “barbed end” than their “pointed end” (Woodrum, Rich et al. 1975) (see Fig. S1), and how does dissociation of the γ-phosphate change the properties of the filament?

Holmes et al. (Holmes, Popp et al. 1990) based the first atomic model of the actin filament on their crystal structure of the actin molecule and x-ray fiber diffraction data. Oda et al. improved the model using X-ray fiber diffraction data to 3.3 Å resolution in the radial direction and 5.6 Å along the equator (Oda, Iwasa et al. 2009) and discovered that the subunits in filaments are flattened compared with monomers. Fujii et al. (Fujii, Iwane et al. 2010) followed with the first cryo-EM structure of the Mg^2+^-actin filament, achieving 6.6 Å resolution before the advent of direct electron detectors. Since then, improvements in cryo-EM methods (von der Ecken, Müller et al. 2014, Galkin, Orlova et al. 2015, von der Ecken, Heissler et al. 2016) have extended the resolution of filament reconstructions to 3.7 Å (Table 1) but have not answered the historic questions owing to limited resolution and the absence of structures with bound ATP. A high-resolution structure (3.4 Å) a filament of bacterial actin, AlfA, (Szewczak-Harris and Löwe 2018) is left handed and lacks subdomain 4 but offers interesting comparisons with actin filaments.

**Table 1.** Comparison of actin filament structures. Note: (—) coordinate files are not available at RCSB PDB.

| Sample (source) | Method, resolution | Rise, Å | Twist, ° | Reference |
| --- | --- | --- | --- | --- |
| Ca-ADP-actin (muscle) with gelsolin | X-ray fiber diffraction, 3.3 Å (radial) 5.6 Å (equatorial) | --- | --- | (Oda, Iwasa et al. 2009) |
| Mg-ADP-actin (muscle) | Cryo-EM, energy filter, CCD camera, 6.6 Å | 27.60 | -166.66 | (Fujii, Iwane et al. 2010) |
| Mg-ADP-actin (muscle) with coronin | Cryo-EM, CCD camera, 8.6 Å | --- | -165.9 | (Ge, Durer et al. 2014) |
| Mg-ADP-BeFx-actin (muscle) with coronin | Cryo-EM, CCD camera, 8.6 Å | --- | -166.3 | (Ge, Durer et al. 2014) |
| ADP-actin (muscle) (Ca-actin in 1 mM MgCl<sub>2</sub>) | Cryo-EM, direct detector, 4.7 Å | 27.63 | -166.68 | (Galkin, Orlova et al. 2015) |
| ADP-actin (muscle) with tropomyosin (Ca-actin in 2 mM MgCl<sub>2</sub>) | Cryo-EM, direct detector, drift correction, 3.7 Å for actin; 3.6 Å in 2016 | 27.44<br>27.20 | -166.40<br>-166.42 | (von der Ecken, Müller et al. 2014, von der Ecken, Heissler et al. 2016) |
| ADP-actin (human $\gamma$) with tropomyosin, myosin head; (Ca-actin in 2 mM MgCl<sub>2</sub>) | Cryo-EM, drift correction, 3.9 Å overall, 3.7 Å for actin | 27.32 | -166.58 | (von der Ecken, Heissler et al. 2016) |
| Mg-AMP-PNP-actin (muscle) | Cryo-EM, direct detector, energy filter, phase plate, drift correction, 3.4 Å | 27.40 | -166.48 | Present work |
| Mg-ADP-P<sub>i</sub>-actin (muscle) | Cryo-EM, direct detector, energy filter, phase plate, drift correction, 3.4 Å | 27.33 | -166.53 | Present work |
| Mg-ADP-actin (muscle) | Cryo-EM, direct detector, energy filter, drift correction, 3.6 Å | 27.52 | -166.63 | Present work |

Here we report high resolution, cryo-EM structures of actin filaments with bound ATP analog β,γ-imidoadenosine 5’-triphosphate (AMPPNP) (3.4 Å), ADP with inorganic phosphate (ADP-P_i_) (3.4 Å) or ADP (3.6 Å), the three well characterized nucleotide states of actin monomers and filaments (Pollard 1986, Fujiwara, Vavylonis et al. 2007, Courtemanche and Pollard 2013). These structures provide insights regarding the different rates of elongation at the two ends of filaments, the timing of the conformational changes associated with polymerization, the mechanism of ATP hydrolysis, and the effects of γ-phosphate dissociation on the filament.

## Results

### Cryo-EM reconstructions of AMPPNP-, ADP-P_i_- and ADP-actin filaments

We obtained near atomic resolution reconstructions of actin filaments with three different bound nucleotides, AMPPNP, ADP-P_i_ or ADP (Fig. 1). To assure that these samples were homogeneous, we purified Ca-ATP-actin monomers from chicken skeletal muscle (which has the same sequence as the more frequently used rabbit skeletal muscle actin) and then converted these monomers to the desired nucleotide states before polymerization. We prepared Mg-ADP-actin monomers by exchanging Mg^2+^ for Ca^2+^ and then treating the Mg-ATP-actin with hexokinase and glucose to make Mg-ADP-actin (Pollard 1984) for polymerization in KMEI buffer (100 mM KCl; 1 mM MgCl_2_; 1 mM EGTA; 10 mM imidazole, pH 7.0) to make Mg-ADP-actin filaments or in KMEI buffer with 10 mM potassium phosphate to make Mg-ADP-P_i_-actin filaments. These precautions make more homogeneous filaments than allowing ATP-actin to hydrolyze the bound ATP and dissociate the γ-phosphate. Since ATP hydrolysis is rapid (Blanchoin and Pollard 2002) and irreversible (Carlier and Pantaloni 1986) in filaments, we used the slowly hydrolyzed ATP analog AMPPNP with a nitrogen atom replacing the oxygen atom bridging β-phosphate and γ-phosphate. AMPPNP-actin polymerizes with the same kinetics as ATP-actin (Courtemanche and Pollard 2013). We exchanged AMP-PNP for ATP to make AMPPNP-actin monomers, which we polymerized in KMEI. We refined methods to make thin films of these filaments on holey, carbon-coated grids for rapid freezing.

**Figure 1.**
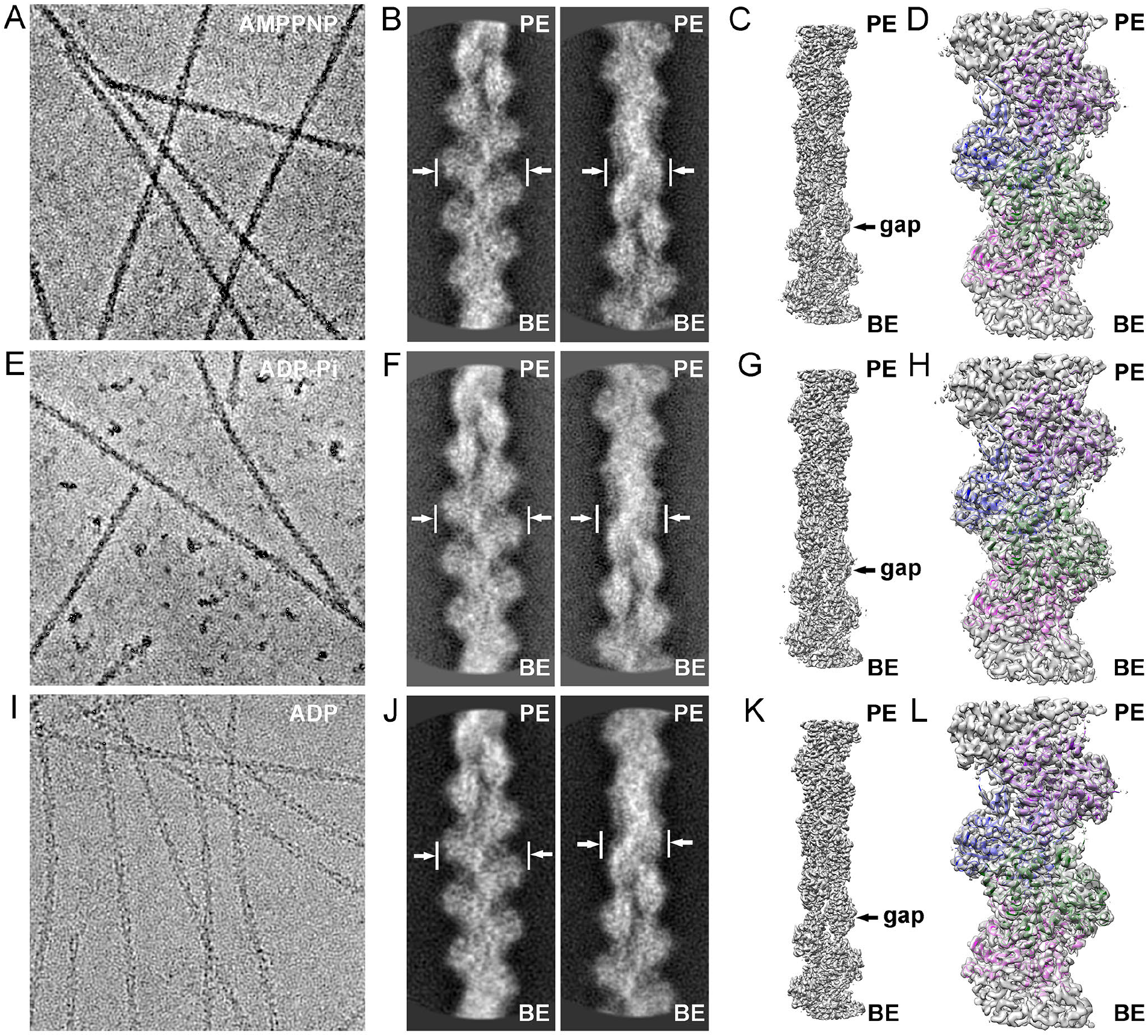
Helical reconstructions of actin filaments from low dose electron cryo-micrographs. (A-D) AMPPNP-actin filaments reconstructed at 3.4 Å resolution. (E-H) ADP-Pi-actin filaments reconstructed at 3.4 Å resolution. (I-L) ADP-actin filaments reconstructed at 3.6 Å resolution. Methods: AMPPNP- and ADP-P_i_-actin filaments were imaged with Volta phase plate and the maps are contoured at 0.030 e/Å^3^. ADP-actin filaments were imaged without the phase plate and the map is contoured at 0.042 e/Å^3^. The pointed end (PE; facing up) and barbed end (BE; facing down) are labeled. (A, E, I) Representative images. (B, F, J) Pairs of contrast-inverted 2D class averages showing the wide and narrow projections of filaments. (C, G, K) 3D reconstructions. (D, H, L) Models fit into the density maps with the chain in each subunit colored differently.

We employed cryogenic electron microscopy (cryo-EM) to collect high-quality images of actin filaments with the three different bound nucleotides (Figs. 1 A, E and I) using a direct-electron detector after an energy-filter. We used a Volta phase plate for imaging AMPPNP- and ADP-P_i_-actin filaments but not the ADP-actin filaments. We selected images with maximum resolution better than 3.8 Å for further analysis. After drift correction and dose weighting, regions from the middles of filaments were boxed out and windowed into segments (particles) for reference-free 2D classifications (Figs. 1B, F and J) and 3D reconstructions (Figs. 1C, G and K) using the iterative helical real-space reconstruction (IHRSR) method (Egelman 2000). The reconstructions were refined to global resolutions of 3.4 Å for AMPPNP-actin filaments, 3.4 Å for ADP-P_i_-actin filaments and 3.6 Å for ADP-actin filaments (Figs. S2 A, C and E) estimated by Fourier shell correlation with 0.143 criteria (FSC_0.143_). We confirmed global resolutions with layer-line images calculated from back projected images (Figs. S2B, D and F) and by calculating local resolution estimations (Fig. S3A, B and C).

The cryo-EM maps clearly resolved the bound nucleotides and side chains of most residues (Figs. 2A, B), allowing us to build atomic models (Figs. 1D, H and L and 2). The only residues missing from the maps were flexible regions at the N-terminus (residues 1-3) and in the DNase I-binding loop (residues 46-48).

### Conformations of actin subunits in filaments with three different bound nucleotides

The backbone conformations of polymerized actin subunits with bound AMPPNP, ADP-P_i_ and ADP are almost identical (Fig. 2C), so assembly, rather than ATP hydrolysis or phosphate release, is responsible for the conformational changes known to occur during actin polymerization (Oda, Iwasa et al. 2009, Fujii, Iwane et al. 2010, Murakami, Yasunaga et al. 2010, von der Ecken, Müller et al. 2014, Galkin, Orlova et al. 2015). The root mean square deviations between the α-carbon atoms in our three structures were <0.5 Å (Supplemental Fig. S4) The only meaningful small difference is in the P1 loop at S14 (see the discussion below and Fig. S8). The conformations of polymerized actin subunits all differ from the corresponding nucleotide states of actin monomers packed in crystals (Fig. 2D) as described in detail below.

**Figure 2.**
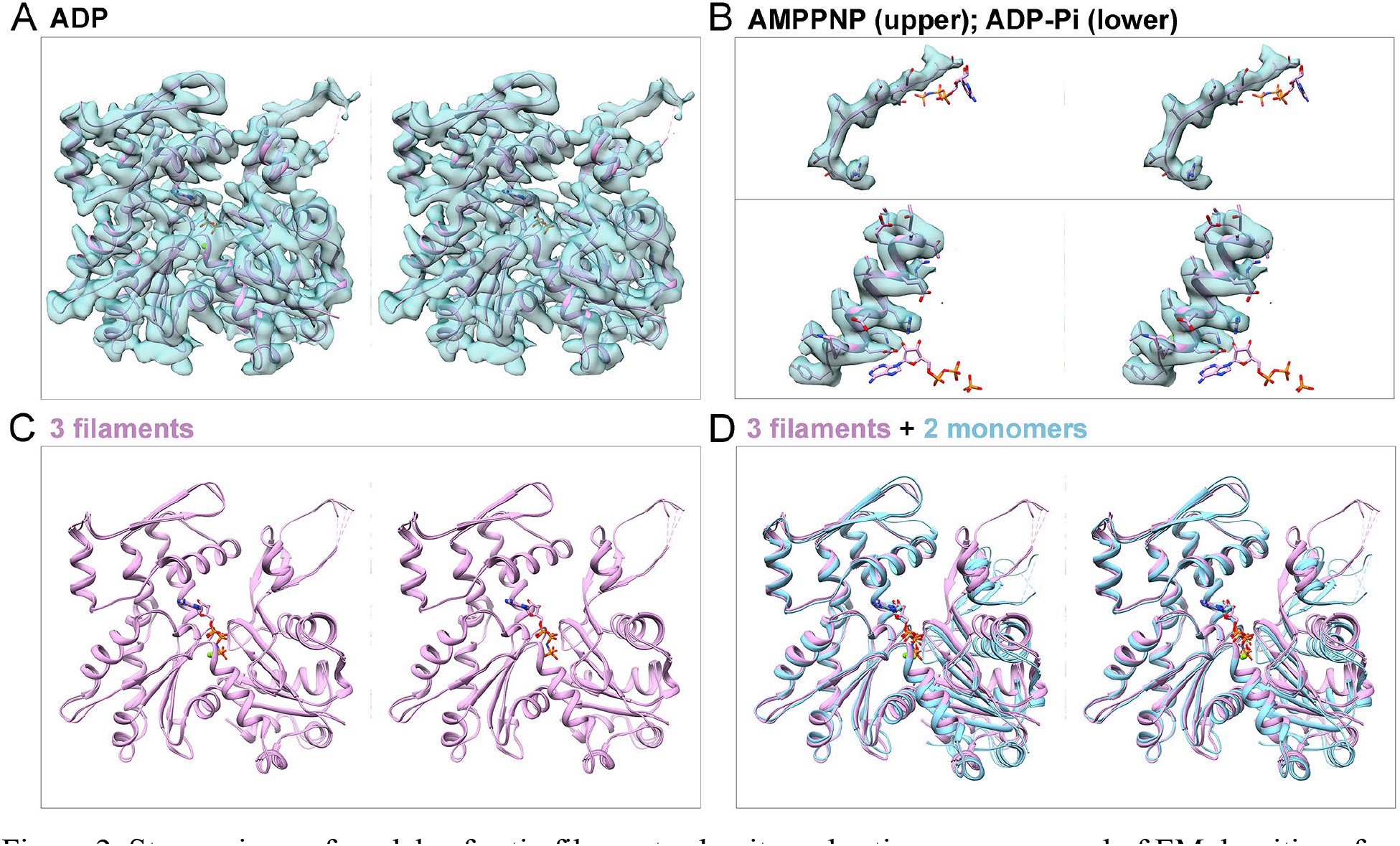
Stereo views of models of actin filament subunits and actin monomers and of EM densities of actin filament subunits contoured at the same levels as Fig. 1. (A) Map and ribbon diagram of one subunit from the ADP-actin filament. (B) Maps and stick figure models show densities for the side chains of a β-strand from the AMPPNP-actin filament (upper) and an α-helix from the ADP-P_1_-actin filament. The stick models of the nucleotides are for orientation. (C) Superimposed ribbon diagrams of subunits from the AMPPNP-, ADP-P_i_- and ADP-actin filaments show that their backbones are nearly identical. (D) Three ribbon diagrams from (C) are superimposed on ribbon diagrams of actin monomers (light blue) with bound Mg-ATP (PDB: 1nm1) or Mg-ADP (PDB: 3a5l) to show differences between filament subunits and monomers. Structures in C and D are aligned using subdomain 3.

The three filament structures have slightly different helical parameters (Table 1 and Fig. S4), the rise and twist (Fig. S1B and C). The three datasets were acquired using the same microscope and camera, so they should be comparable. The rise per subunit for ADP-actin is 0.12 Å (0.4%) larger than AMPPNP-actin and 0.19 Å (0.7 %) larger than ADP-P_i_-actin, so the filament expands longitudinally a small amount upon ATP hydrolysis and phosphate release.

Most other actin filament structures with sub-nanometer resolutions have bound ADP, all based on cryo-EM data except for one based on x-ray fiber diffraction data (Table 1). In one case the filaments were capped with gelsolin (Oda, Iwasa et al. 2009), others had bound tropomyosin (von der Ecken, Müller et al. 2014) or tropomyosin and myosin (von der Ecken, Heissler et al. 2016), one had bound coronin and BeF_x_ (Ge, Durer et al. 2014) and three had mixtures of Ca-ADP- and Mg-ADP-subunits (von der Ecken, Müller et al. 2014, Galkin, Orlova et al. 2015, von der Ecken, Heissler et al. 2016). Only our Mg-ADP-actin filament model and that of Fujii et al. (Fujii, Iwane et al. 2010) (PDB: 3mfp) are the same chemical state. These two Mg-ADP-actin structures have similar helical parameters (Table 1) even though the data were acquired and processed differently. Our ADP-actin model fits reasonably well into Fujii’s lower resolution map (EMDB: 5168), but the subunits in the two models differ substantially (RMSD = 1.62 Å), largely because the nucleotide binding cleft is closed more tightly in our model (Fig. S4C and H). The subunits in filaments of human ADP-α-actin decorated with tropomyosin are close to our model of ADP-actin (RMSD: 0.65 Å; Fig. S4D and I), but the rise per subunit in the decorated filament is smaller than all of our models, so the bound proteins may make actin filaments more compact.

### Conformational changes within subunits associated with polymerization

Actin subunits are flattened to the same extent in filaments with bound Mg-AMPPNP, Mg-ADP-P_i_ or Mg-ADP (Figs. 2D, 3 and S4F, G). The centers-of-mass (COM) of the four subdomains form a dihedral angle-like structure (Fig. S5A). The inter-domain dihedral angle of subunits in our Mg-AMPPNP-actin filament model are rotated 13.1° relative to the crystal structure of the Ca-ATP-actin monomer bound to DNase-I (PDB: 2a42) (Fig. 3A) and by 18.7° relative to the TMR-labeled Ca-ADP-actin monomer (PDB: 1j6z) (Fig. S6). DynDom analysis (Hayward and Lee confirmed that rotation occurs around a hinge helix (residues 137-145) and a hinge loop (residues 335-337) as described previously (Oda, Iwasa et al. 2009, Fujii, Iwane et al. 2010). The catalytic residue Q137 stands on the hinge helix. The side chain of K336 in the center of the hinge loop interacts with the adenosine base as in monomers (Kabsch, Mannherz et al. 1990).

**Figure 3.**
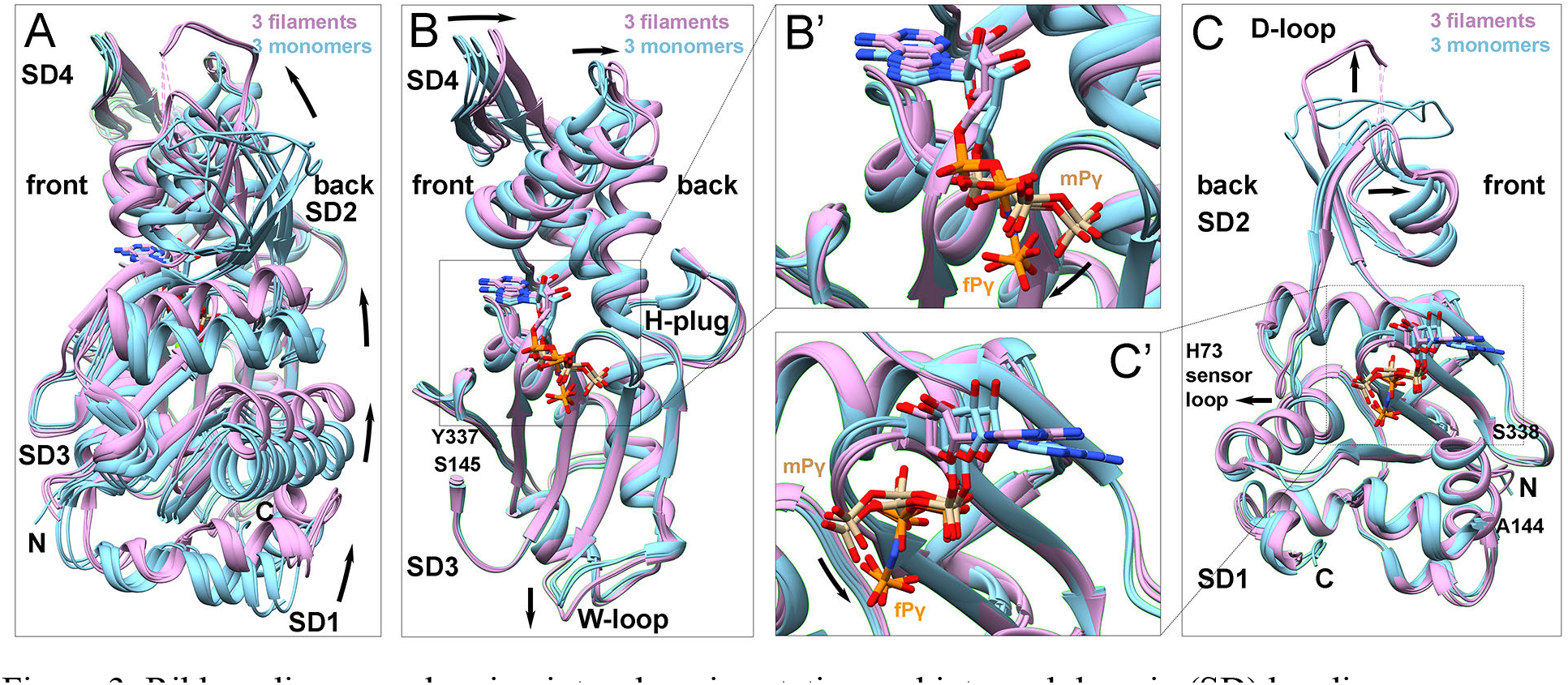
Ribbon diagrams showing inter-domain rotation and inter-subdomain (SD) bending upon filament formation. (Light blue) three actin monomer crystal structures in the closed conformation: rabbit skeletal muscle Ca-ATP-actin complexed with DNase I (PDB: 2a42); *Dictyostelium* Mg-ADP-actin complexed with human gelsolin segment 1 (PDB: 3a5l); and budding yeast Mg-ATP-actin complexed with human gelsolin segment 1 (PDB: 1yag). (Plum) Our three actin filament EM structures in the tightly closed conformation: AMPPNP-, ADP-P_i_- and ADP-actin. Nucleotides are shown as stick figures with phosphorus atoms orange in filaments and tan in monomers. The γ-phosphate group in filaments (fP_γ_) points downward compared with monomers (mP_γ_). Arrows mark differences between monomers and filaments. (A) Dihedral angle-like inter-domain rotation. The six molecules are aligned using subdomains 2 and 4 (residues 145-337). (B) Bending of subdomains 3 (residues 145-180 and 270-337) and 4 (181269). The molecules are aligned using subdomain 3. Inset B’ shows the difference between the phosphates in monomers and filaments. (C) Bending of subdomains 1 and 2. The molecules are aligned using subdomain 1 (residues 5-32, 70-144, and 338-370). Inset C’ shows the difference between the phosphates in monomers and filaments.

We confirm the discovery of Fujii, et al (Fujii, Iwane et al. 2010) that the subdomains in both halves of actin also flatten during polymerization (Fig. 3B, C). Subdomains 3 and 4 are flatter in filaments than monomers as subdomain 4 tilts toward the back side of actin relative to subdomain 3 (Fig. 3B). A subtle movement tilts subdomain 2 toward the front side of actin relative to subdomain 1 (Fig. 3C) as the D-loop (residues 40-50) moves upward into a more extended conformation.

### Contacts between subunits in filaments

Each subunit in the middle of a filament has four neighbors making contacts that together bury 3540 Å^2^ of surface area (Fig. S1B). The “interstrand” interactions along the short pitch helix (subunits *m+1* to *m* to *m-1*) (Fig. 4; see Fig. S1B) are weaker than the “intra-strand” interactions along the long-pitch helix (subunits *m+2* to *m* to *m-2*) (Fig. 5). Each “inter-strand” contact buries 490 Å^2^ of surface area, so the contacts with *m+1* and *m-1* bury a total of 980 Å^2^ of surface area on subunit m. Along the long-pitch helix interactions of subdomains 2 and 4 of each subunit with subdomain 3 of the neighbor toward the pointed end bury 1180 Å^2^ of surface area for a total of 2360 Å^2^ with both neighbors. The stronger contacts along the long-pitch helix likely explain why longitudinal dimers are favored over short-pitch dimers for the first step in nucleation (Sept and McCammon 2001).

**Fig. 4.**
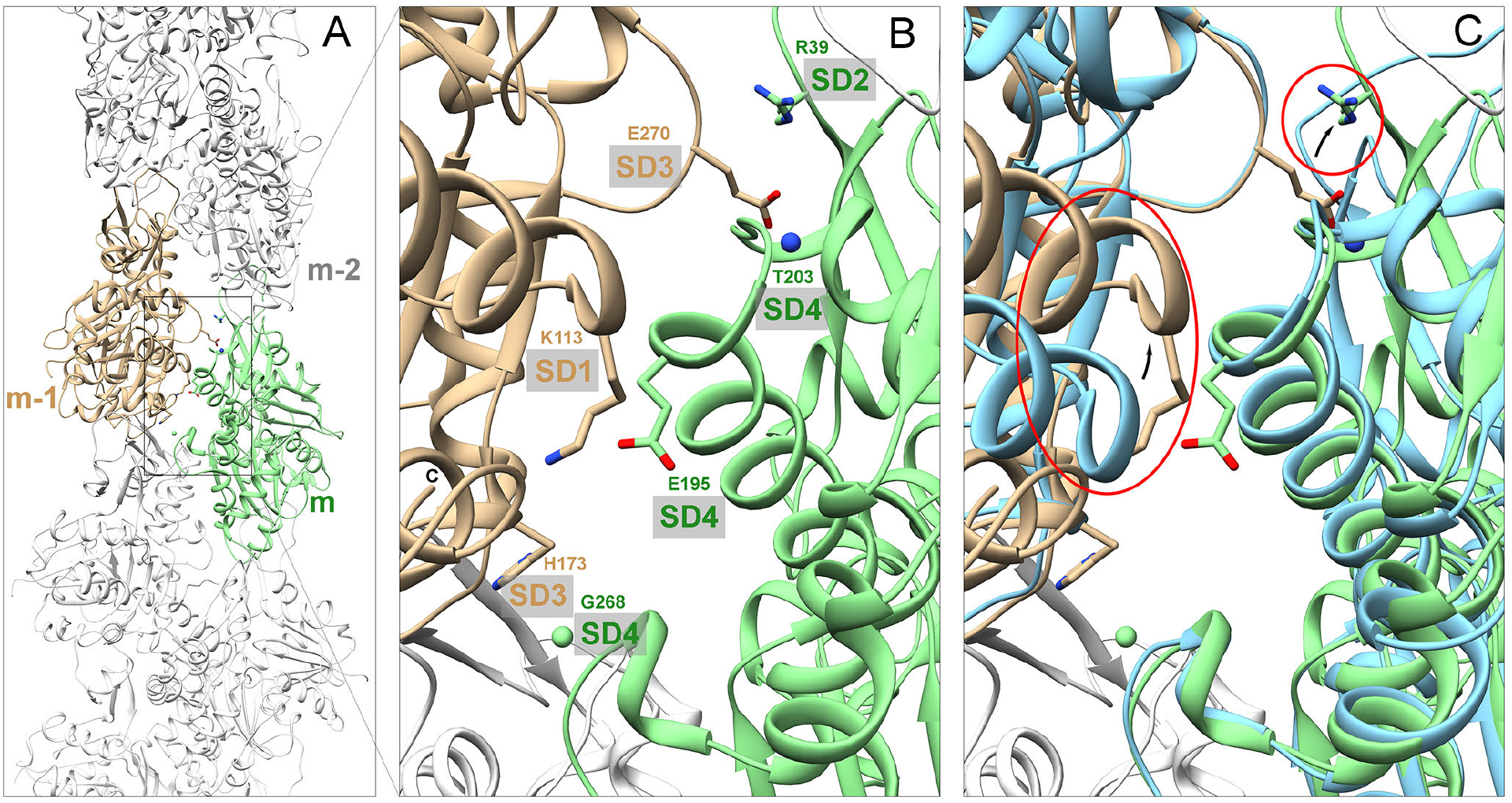
Ribbon diagrams showing lateral contacts between subunits along the short-pitch helix. (A) Overview of filament with two subunits highlighted in green and tan. (B) Detail of contacts between subunits *m* (green) and *m-1* (tan) with labels on stick figures of the interacting residues. (C) Same as (B) with two overlaid actin monomers (light blue) aligned with subdomains 3 and 4. This superimposition shows that subunit flattening allows interactions between the backbone and side chain of E195 in subdomain 4 of subunit *m* and K113 in subdomain 1 of subunit *m-1* and between side chain of E270 in subdomain 3 of subunit *m-1* with side chain of R39 in subdomain 2 of subunit *m*

**Figure 5.**
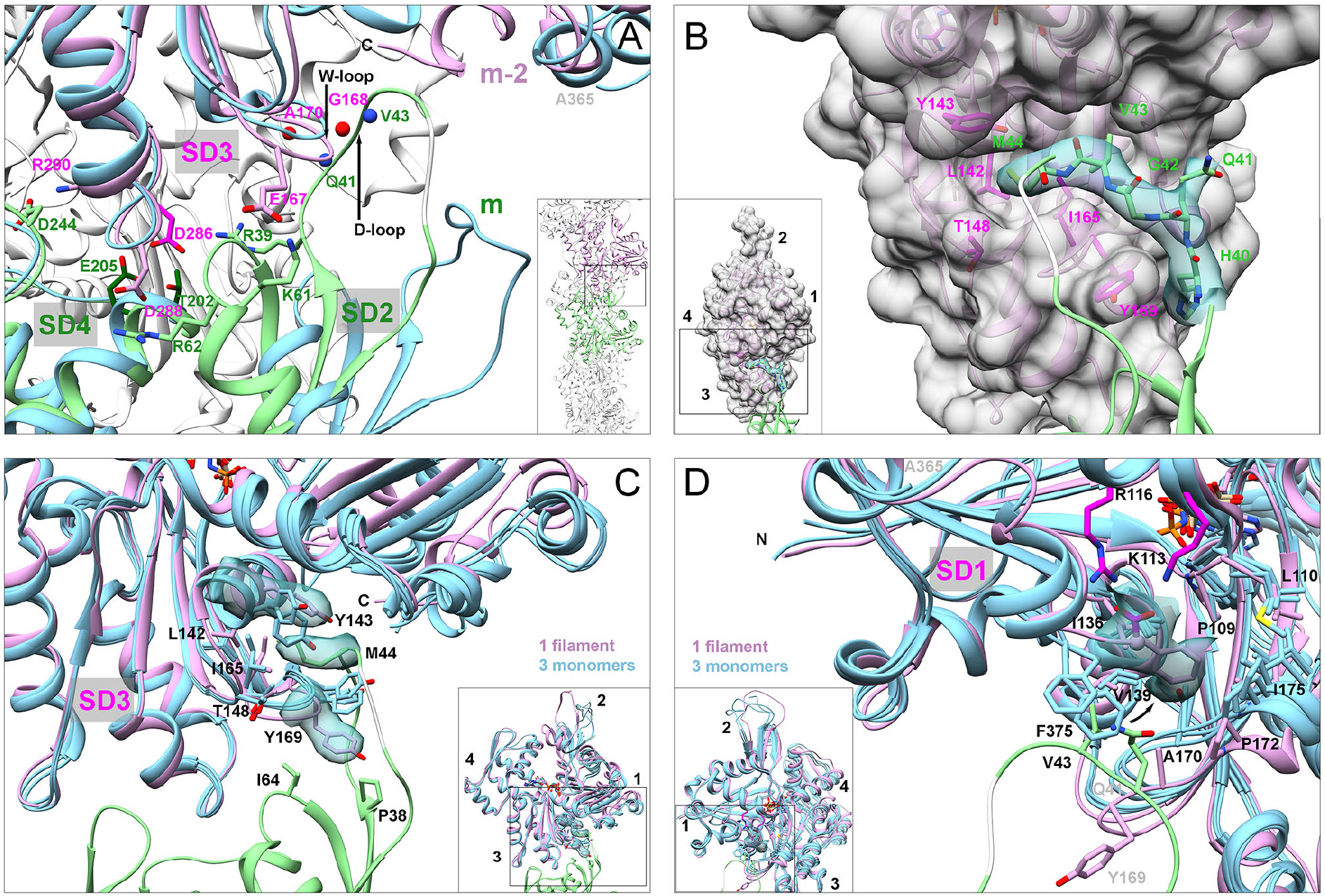
Interactions along the long-pitch helix of the AMPPNP actin filament between subdomains 2 and 4 of subunit *m* (green) and subdomain 3 of subunit *m-2* (plum). Ribbon diagrams show the backbones, stick figures show key side chains, semi-transparent turquoise maps at same contour levels as Fig. 1 show some experimental densities and grey surface in (B) is calculated from the model. The conformational changes that enable these interactions are illustrated by superimposing ribbon diagrams (light blue) of actin monomers aligned on subdomains 3 and 4 in (A), subdomain 3 in (C) and subdomain 1 in (D). The monomer structures are Ca-ATP-actin complexed with DNase I (PDB: 2a42; in A, C and D), Mg-ADP-actin with human gelsolin segment 1 (PDB: 3a5l; in C and D) and Mg-ATP-actin with human gelsolin segment 1 (PDB: 1yag; in C and D). (A) Polar contacts illustrated by stick figures of interacting side chains: R39 in subdomain 2 with D286 in subdomain 3; K61 in subdomain 2 with E167 in subdomain 3; R62 in subdomain 2 with D288 in subdomain 3; and D244 in subdomain 4 with R290 in subdomain 3. The side chains of T202, E205 and D286 form the binding site for the “polymerization cation”. (B) The D-loop (residues 40-50) of subunit *m* wraps snugly around the W-loop (residues 165172) of subunit *m-2.* The density for residues 46-48 is weak, so only the position of the backbone is shown in light grey. (C) Downward bending of the W-loop with Y169 at its tip (density from the AMPPNP-actin filament) in filaments separates Y143 from Y169 and opens a hydrophobic pocket for M44 from subunit *m*. The M44 side chain from the D-loop (density from the AMPPNP-actin filament) of subunit *m* inserts into a hydrophobic pocket in subunit *m-2* formed by L142, Y143, T148 and I165 above the W-loop. (D) Conformations of the C-terminal F375 vary in crystal structures of actin monomers but in all three filament structures its side chain is buried in a hydrophobic pocket formed by P109, L110, I136, V139, A170, P172 and I175 of its subunit and V43 of the lower subunit (*m*). The C-terminal carboxyl group (magenta) is turned outward close to the magenta side chains of K113 and R116 in subdomain 1 its own subunit.

### Conformational changes on the surface of subunits associated with polymerization

Our structures show that local conformational changes during polymerization create complementary surfaces for interactions between the subunits along both the short-pitch (Fig. 4) and long-pitch helices (Fig. 5). These interfaces are virtually identical in filaments with each of the bound nucleotides, so we describe them together in Fig. 5. The contacts in our three models are similar to those in the model of Ca/Mg-ADP-actin-tropomyosin-myosin filaments (von der Ecken, Heissler et al. 2016).

*Interactions along the short-pitch helix:* Inter-strand contacts are largely (~87%) between subdomain 3 at the barbed end of one subunit and subdomain 4 at the pointed end of its neighbor (Fig. 4). Subunit flattening is required for three polar contacts between the subunits: (i and ii) a hydrogen bond between the backbones of K113 of subunit *m-1* and E195 of subunit *m* and charge-charge interaction of their side chains; and (iii) electrostatic interaction of the side chain of E270 in subdomain 3 of subunit *m-1* and side chain of R39 in subdomain 2 of subunit *m*. Subunit flattening is not required for two other contacts: a hydrogen bond between the side chain of E270 (in H-plug) with the backbone of T203 in the upper part of the nucleotide-binding cleft formed by subdomains 2 and 4 of subunit *m*; and a contact between the backbone of G268 in the H-plug of subunit *m* with the sidechain of H173 in subdomain 3 of subunit *m-1*.

*Interactions along the long-pitch helix:* Longitudinal contacts involve interactions of subdomains 2 and 4 of subunit *m* with subdomain 3 at the barbed end of subunit *m-2* (Fig. 5). Subdomain 2 is small, but makes 3 pairs of charge-charge interactions, 2 pairs of backbone hydrogen bonds and large hydrophobic contacts with its neighbor.

Subdomains 2 and 4 make multiple polar contacts with the barbed end subdomain 3 of subunit *m-2* (Fig. 5A). The side chains of T202, E205 in subdomain 4 of subunit *m* and D286 in subdomain 3 of subunit *m-2* are proposed to bind the “polymerization cation” that stabilizes the filament (Kang, Bradley et al. 2013). Our maps have weak density in the position proposed for the polymerization site cation; the densities of these acidic side chains are also weak, as usual in EM maps (Wang and Moore 2017). Another divalent cation bound to subdomain 2 is proposed to stiffen the filament (Kang, Bradley et al. 2013), but none of our maps has density at the position proposed for this cation. However, the stiffness-related cation ion might be located between the side chain of D56 in subdomain 2 and the backbone of V30 in subdomain 1, as seen in the crystal structure of TMR-labeled actin monomer (Otterbein, Graceffa et al. 2001)(PDB: 1j6z).

Interaction of the D-loop (residues 40-50) of subunit *m* with subdomain 3 of subunit *m-2* forms a major contact along the long-pitch helix (Figs. 5). The D-loop is flexible in monomers, so the electron densities are incomplete or weak in most crystal structures unless bound to associated proteins. All three of our maps of filaments have densities for D-loop residues 40-44 (including the side chain of M44), which are immobilized in filaments. The density for V45, G46, M47 and G48 in our maps and other maps (von der Ecken, Heissler et al. 2016) (PDB: 5jlf) is weak, so these residues are flexible or have multiple conformations. We only include a provisional model for the backbone of these residues.

Our structures reveal that the interactions of the D-loop with subdomain 3 of the adjacent subunit depend on displacement of the W-loop of subdomain 3 (residues 165-172, named for its interactions with WH2 motifs) from its position in monomers towards the barbed end of the subunit (Figs. 3B and 5B, C). The α-carbon of Y169 at the tip of the W-loop moves about 2.2 Å. The backbones of G168 and Y169 in our three models bend toward the barbed end and are similar to the 3.7 Å model of the ADP-actin-tropomyosin filament (von der Ecken, Müller et al. 2014) (PDB: 3j8a) but both differ from the updated 2016 model of the ADP-actin-tropomyosin filament (von der Ecken, Heissler et al. 2016) (PDB: 5jlf). These structures alone do not explain how flattening of the subunits in the filament drives this conformational change in the W-loop, but this small change does facilitate two crucial interactions of the D-loop of subunit *m* with subunit *m-2.*

First, movement of the W-loop allows the D-loop of subunit *m* to wrap around the W-loop (Fig. 5 B and C). The side chain of Y169 (F169 in yeast) has multiple rotamer conformations in monomers (Fig. 5C), but hydrophobic contacts with P38 and I64 of subunit *m* immobilize the W-loop in a single conformation in all three of our filaments (Fig. 5C). Molecular dynamics simulations by Zheng, et al. (Zheng, Diraviyam et al. 2007) suggested the importance of Y169 in filament formation, which was confirmed by biochemical experiments (Kudryashov, Grintsevich et al. 2010).

Second, movement of the W-loop opens a hydrophobic pocket surrounded by L142, Y143, T148 and I165 into which the side chain of M44 from the D-loop of subunit *m* inserts. This contact buries 151 Å^2^ of surface area. Von der Ecken et al. (von der Ecken, Müller et al. 2014) described M44 in this hydrophobic pocket but not that the pocket in monomers is too small to accommodate the M44 side chain. The distance between the Y143 CZ atom (next to the hydroxyl group) and the a-carbon of Y169 increases from 6.6 Å in monomers to 10.9 Å in filaments, avoiding a clash of the M44 side chain with the aromatic ring of Y143 and backbone of Y169.

The conformation of the C-terminal tail is disordered in actin monomers but adopts the same folded conformation in all three of our filament structures. The side chain of C-terminal F375 is buried in a large hydrophobic pocket formed by residues from subdomain 1 (P109, L110, I136, V139) and subdomain 3 (A170, P172, I175) of the same subunit, and V43 from the D-loop of subunit *m+2*. The C-terminal carboxyl group is close enough to the side chains of K113 and R116 for favorable electrostatic interactions (Fig. 5D). On the other hand, the map of the crystal of DNase I-Ca^2+^-ATP-actin (PDB: 2a42) has no density for residues 366-375 (Fig. 5D) and the densities for the C-termini in other crystals (PDBs: 3a5l and 1yag) do not superimpose well (Fig. 5D). These various conformations of the side chain of F375 in monomers may interfere with the association of the D-loop of subunit *m+2* due to steric clashes with the side chain of Q41 in the D-loop (Fig. 5D).

### Interactions of the three nucleotides with actin subunits in filaments

Our three high-resolution maps and models of actin filaments (Figs. 6, 7 and S7) show that polymerization does not disrupt any of the interactions between the protein and base or ribose of ATP known from crystal structures of ATP-actin and ATP-Arp2/3 complex (Kabsch, Mannherz et al. 1990, Vorobiev, Strokopytov et al. 2003, Nolen and Pollard 2007, Dominguez and Holmes 2011). However, the active site of AMPPNP-actin filaments differs in several functionally important ways from crystal structures of ATP-actin monomers.

First, all three phosphates are displaced toward the barbed end of the subunit during polymerization (Figs. 3 and 7), with the α- and β-phosphates moving 1.0 Å and the γ-phosphate 3.0 Å concomitant with the loss of hydrogen bonds between non-bonded oxygens of the γ-phosphate and backbone nitrogens of D157, G158 and G159 in subdomain 3. Subunit flattening moves the backbone of S14 and G15 ~1.8 Å toward the front side of the subunit and reduces the length of the hydrogen bond between backbone nitrogen of S14 and the β-phosphate from ~3.5 Å in monomers to ~2.5 Å in filaments (Fig. S8), changes that probably contribute to the movement of the phosphates.

Second, rotation of the whole outer domain with respect to the inner domain repositions Q137 on the hinge helix of subdomain 1 (Figs. 7 and S7). This brings the side chain OE1 atom of Q137 2.2 Å closer to the γ-phosphorous atom, which are separated 5.4 Å in monomers and 3.2 Å in filaments.

Third, the rotamer configuration of the H161 side chain in all three of our filaments brings the side chain NE2 atom ~2.5 Å closer to the γ-phosphorous atom than in most monomers (9.1 Å) (Figs. 7A, C, D and S7). An exception is the crystal structure of the *Dictyostelium* Li-ATP-actin monomer bound to human gelsolin subdomain 1 (Vorobiev, Strokopytov et al. 2002) (PDB: 1nmd), where the side chain of H161 is in a conformation between these two rotamers, 7.8 Å from the γ-phosphorous atom (Fig. 7B).

### Changes in filaments associated with ATP hydrolysis and phosphate release

Conformational changes in actin subunits during polymerization (Fig. 3) reposition the side chains of Q137 and H161 relative to the γ-phosphate and promote hydrolysis (Fig. 7), but the overall structures of the subunits do not change upon ATP hydrolysis and phosphate release (Figs. 2C and 6). The densities corresponding to the γ-phosphate differ in our three maps.

The reconstructions of the AMPPNP- and ADP-P_i_-actin filaments are remarkable similar, both having strong density for the γ-phosphate (Fig. 6A, C). The only significant change after hydrolysis of ATP is the position of the γ-phosphate, which moves to a position midway between the β-phosphate and the side chain of Q137 (Fig. 6A, C). When contoured at the same level, the densities for the γ-phosphates are similar in the maps of AMPPNP-actin and ADP-P_i_-actin, so the γ-phosphate site is fully occupied. The positions of backbone and all the side chains in the active site are the same in AMPPNP- and ADP-P_i_-actin filaments.

**Figure 6.**
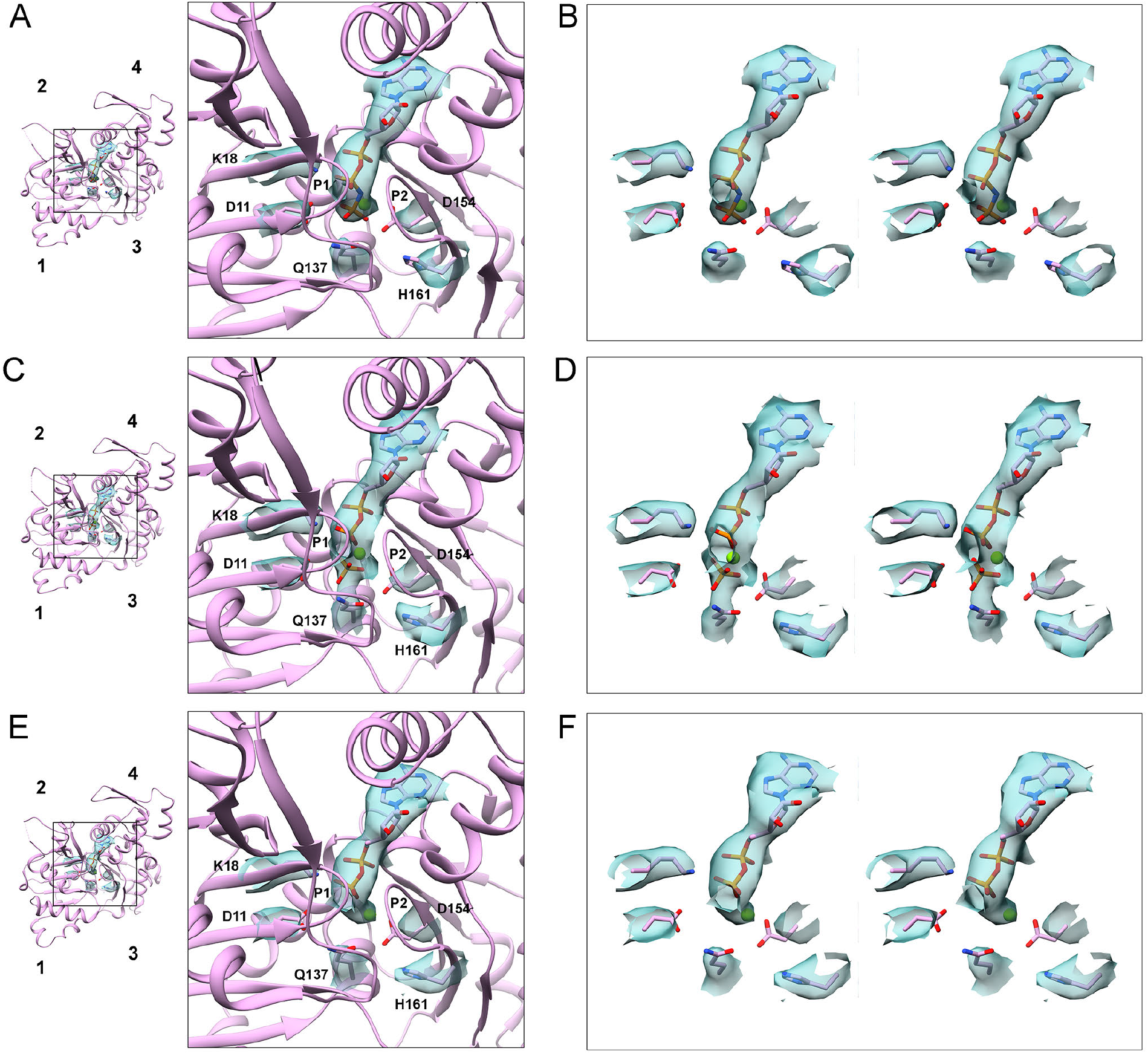
Changes in the active site during the ATPase cycle of polymerized actin. (A, B). AMPPNP-actin. (C, D) ADP-P_i_-actin. (E, F) ADP-actin. Each row has three parts: (left) a small ribbon diagram looking down into the active site for orientation; (center) a ribbon diagram with stick figures and map densities zoned within 2.12 Å of the nucleotide and the important side chains D11, K18, Q137, D154 and H161; and (right) a wall-eye stereo view of the active site densities and stick figures of important side chains.

**Figure 7.**
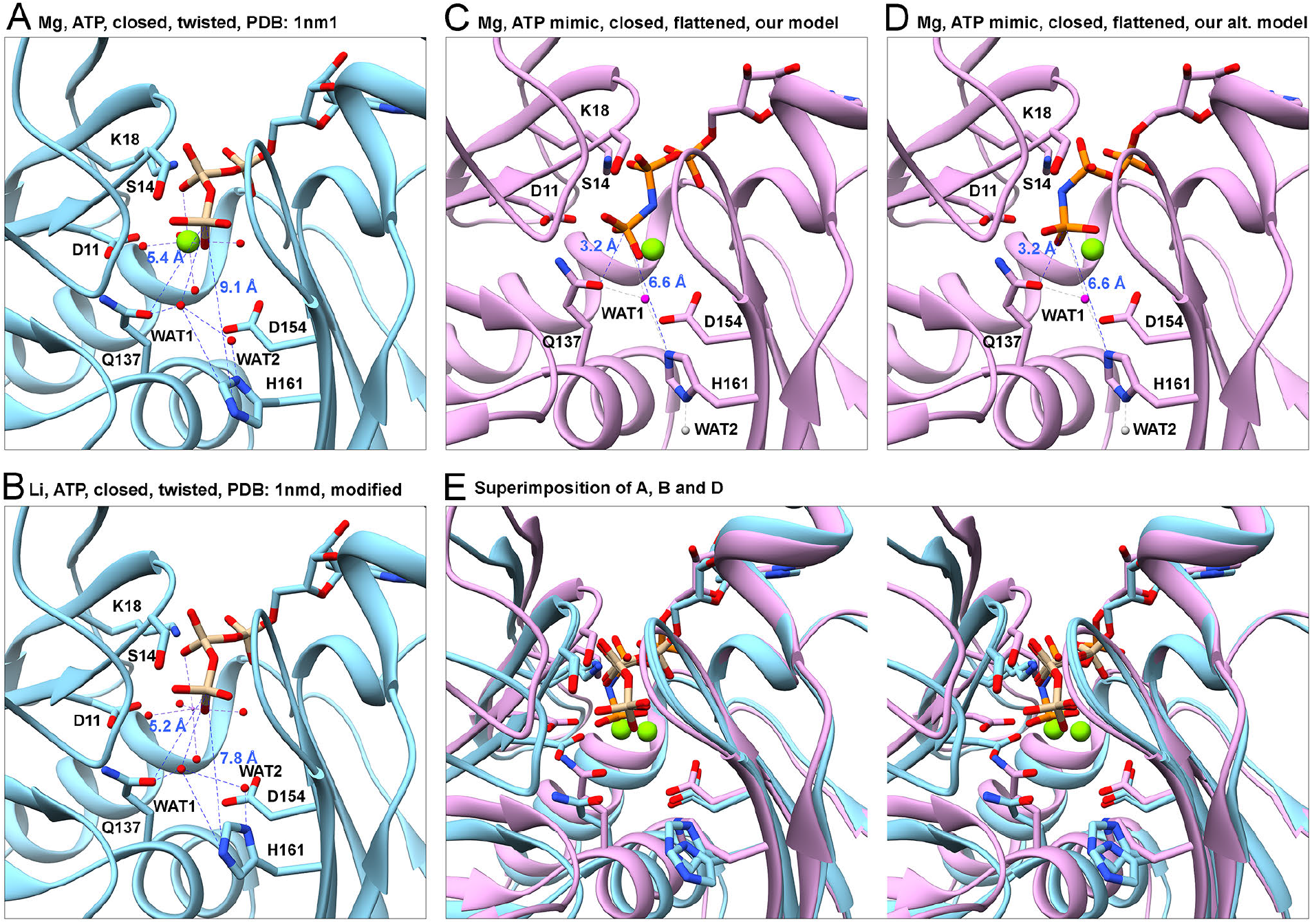
Rearrangement of the catalytic center stimulates ATP hydrolysis by polymerized actin. Ribbon models with stick figures of the nucleotides and selected side chains compare distances between the γ-phosphate (Pγ) and the OE1 atom of Q137, NE2 atom of H161 and water molecules in actin monomers and filaments. (A) Crystal structure at 1.8 Å resolution of the *Dictyostelium* Mg-ATP-actin monomer complexed with human gelsolin segment 1 (PDB: 1nm1). (B) Crystal structure at 1.9 Å resolution of the *Dictyostelium* Li-ATP-actin monomer complexed with human gelsolin segment 1 (PDB: 1nmd). Based on an analysis of the water network, we flipped the imidazole ring of H161, which fits in the electron density equally well as its conformation in the original PDB file. This flipping brings the side chain ND1 atom closer to WAT2 (distance: 2.5 Å) indicating that they are hydrogen bonded. (C) Our model of polymerized Mg-AMPPNP-actin. Compared with monomers, P_γ_ is 2.2 Å closer to the OE1 of Q137 and 2.5 Å closer to the NE2 atom of H161, which is rotated relative to monomers by ~120° to the rotamer observed in filaments in all three nucleotide states (Fig. 6). Water molecules do not appear in the EM map, but we propose that WAT1 from the x-ray structure (magenta ball) remains close to OE1 of Q137 and that WAT2 moves away, either downward (grey ball) or upward (not shown). (D) Our alternative model of polymerized Mg-AMPPNP-actin with a different conformation of α-, β-, γ-phosphates that fits the EM density equally well as (C). The positions of the three phosphorus atoms are almost identical to those in (C), but the positions of bonded oxygen atoms differ. In this conformation, WAT1 is almost in line with γ-phosphorus atom and the bridging atom (N3B in AMPPNP, O3B in ATP) between β- and γ-phosphate, facilitating the attack of WAT1 on the γ-phosphorus atom. (E) Stereo pairs superimposing (A), (B) and (D) aligned on subdomain 3 to show their differences.

The ADP-bound filaments lack density for the γ-phosphate (Fig. 6E, F). Dissociation of phosphate results in the loss of its Mg-O bond with the divalent cation. The only meaningful change in the protein backbone after phosphate dissociation is a small rotation at S14 (Fig. S8). Based on the backbone densities, the α-carbon and side chain of S14 in subdomain 1 adopt a different conformation in the ADP-actin filament than the AMPPNP- and ADP-P_i_-actin filament models. In the AMPPNP- and ADP-P_i_-actin models, the side chain OG atom of S14 is close enough (~3.4 Å) to form a hydrogen bond with the backbone N atom of G74 in the sensor loop of subdomain 2 (residues 71-77) (Fig. S9). In the ADP-actin filament, the side chain of S14 flips towards β-phosphate and the backbone of G158 (in P2 loop of subdomain 3) close enough to make a H-bond with a non-bridging oxygen on β-phosphate. Thus, phosphate dissociation results in a loss of a H-bond between subdomain 1 and the sensor loop, which connects subdomains 1 and 2. Similarly, in actin monomers the side chain of S14 makes H-bonds with a non-bonded oxygen of the γ-phosphate of ATP and the backbone nitrogen of G74 but adopts different rotamers in ADP-actin monomers (Otterbein, Graceffa et al. 2001) (Fig. S8 H and I).

All three filament reconstructions have densities for the divalent magnesium ion (Mg^2+^), rendered as a green ball in Fig. 6. In AMPPNP-actin the protein atom nearest the Mg^2+^ is the side chain OD1 atom of D154 (~3.2 Å). Since an ideal Mg-O bond is ~2.3 Å long, the protein likely contacts Mg^2+^ indirectly through coordinated water molecule as in crystal structures of actin monomers (Vorobiev, Strokopytov et al. 2003). In the absence of the γ-phosphate, the hexa-coordinated magnesium ion remains in the same place, but on the edge of the density that includes the ß-phosphate next to the side chain of D154 (Fig. 6E, F).

The side chain of methylated H73 (Hic73 in the sensor loop) has the same conformation in AMPPNP- and ADP-P_i_-actin filaments but undergoes a small conformational change after phosphate dissociation (Fig. S9). During polymerization, the sensor loop moves toward the backside of actin by 1.1 Å (Fig. 3C). The change after phosphate release is unlikely to impact the interactions between subunits.

## Discussion

### When do conformational changes take place in actin subunits during polymerization, ATP hydrolysis and phosphate release?

Our high-resolution structures establish that the major conformational changes take place when ATP-actin is incorporated into a filament rather than being associated with ATP hydrolysis or phosphate release. The conformational changes during phosphate release are remarkably subtle, in spite of impacting subunit dissociation and binding of cofilin and other proteins. This conclusion was not clear previously, since no prior filament structure had bound ATP. Our findings differ from early reports of conformational changes in subdomain 2 associated with phosphate release in low resolution reconstructions actin filaments (Belmont, Orlova et al. 1999) and spectroscopic assays (Moraczewska, Wawro et al. 1999).

### Why does the barbed end elongate faster than the pointed end?

Flattening actin subunits during incorporation into a filament is associated with conformational changes in the nucleotide cleft (Fig. 7) and on the surface of each subunit (Figs. 3, 4 and 5) that facilitate interactions within the filament. First, displacement of the W-loop (Figs. 3B and 5B, C) creates a knob around which the D-loop of the adjacent subunit binds (Fig. 5C). Second, repositioning the W-loop opens a hydrophobic cavity for insertion of the side chain of highly conserved M44 from the neighboring D-loop (Fig. 5B, C). The importance of this interaction is confirmed by the effects of the enzyme MICAL, which stereo-specifically oxidizes both M44 and M47, causing fast filament disassembly (Grintsevich, Ge et al. 2017). Third, rearranging the C-terminus during polymerization eliminates steric interference between the side chain of F375 and the D-loop of the neighboring subunit (Fig. 5D). Fourth, flattening of the polymerized subunit enables two interactions between K113 and E195 and other polar residues along the short-pitch helix (Fig. 4).

The conformational changes associated with polymerization offer an explanation for the different rates of subunit association at the two ends of filaments (Pollard 1986). Incoming subunits at the barbed end of a filament bind to the side of the terminal subunit (*n*) and the barbed end of the penultimate subunit (*n-1*) (Fig. S6B), so the conformations of these two subunits influence the rate of the reaction. We suggest that the surrounding subunits influence the conformations of these two terminal subunits: subunit *n* interacts with subunits *n-1* and *n-2*; and subunit *n-1* interacts with subunits *n, n-2* and *n-3.* These combined interactions are likely to flatten subunits *n* and *n-1* and change the conformation of the side of subunit *n* and the barbed end of subunit *n-1* to create a favorable, filament-like binding site for the incoming subunit. Therefore, at the barbed end of a filament an incoming subunit with a flexible D-loop interacts with two receptive terminal subunits, which is a favorable reaction.

On the other hand, the D-loops of the two subunits exposed at the pointed end of a filament have no lateral or longitudinal interactions (Fig. S6C) and likely remain as flexible as in monomers. The barbed end of an incoming monomer lacks the features required for favorable interactions with a flexible D-loop on the penultimate subunit at the pointed end, so the reaction is unfavorable. In addition, a low-resolution reconstruction of the pointed end suggested that the H-plug and D-loop of the terminal subunit are folded against the penultimate subunit, compromising subunit association (Narita, Oda et al. 2011).

Thus, the favorable conformation of the barbed end of the filament explains why subunit addition is faster there than at the pointed end (Woodrum, Rich et al. 1975). Factors that are not apparent in our structures of the middle of filaments must contribute to why ATP-actin and ADP-P_i_-actin monomers have larger association rate constants than ADP-actin at both ends (Pollard 1986, Fujiwara, Vavylonis et al. 2007, Courtemanche and Pollard 2013).

### How does polymerization increase the rate of ATP hydrolysis?

Flattening of the actin subunit during polymerization reorganizes the active site in filaments and stimulates ATP hydrolysis (Straub and Feuer 1950) by 42,000-fold from 0.000007 s^-1^ by monomers (Rould, Wan et al. 2006) to 0.3 s^-1^ in filaments (Blanchoin and Pollard 2002). Water is important for ATP hydrolysis, but water molecules do not appear in our cryo-EM maps. Multiple factors contribute to the absence of water: the resolution is limited, radiation creates damage during imaging and water molecules and carboxylate groups are underrepresented in electron potential maps produced by electron microscopy compared with electron density maps from X-ray crystallography (Wang and Moore 2017). Therefore, we will have to depend on MD simulations (Saunders and Voth 2011) of filaments and comparisons with crystal structures of monomers to model the water in the active site of polymerized actin.

Two water molecules in high resolution crystal structures of actin monomers may contribute to hydrolysis (Figs. 7A, B). Water 1 is hydrogen-bonded to the OE1 atom of Q137 (distance: ~3.0 Å), and positioned among the OE1 atom of Q137, the imidazole ring of H161 and the γ-phosphate. Water 2 is located between ND1 atom of H161 and Water 1. H161 is the best candidate to activate the attacking water through H-bonds by extracting a proton from water 1 directly or via the bridging Water 2 (Vorobiev, Strokopytov et al. 2003). The geometries of the γ-phosphate, cation, attacking water, Q137 and H161 in high-resolution crystal structures of Ca-ATP-, Mg-ATP- and Li-ATP-actin monomers are consistent with their relative rates of hydrolysis (Vorobiev, Strokopytov et al. 2003). Cardiac actin with the Q137A substitution polymerizes but releases the γ-phosphate 4-fold slower than normal (Iwasa, Maeda et al. 2008). A measurement at a single time point indicated that hydrolysis was rate limiting for phosphate release by the mutant actin.

Previous work showed that the side chain of Q137 is closer to the nucleotide in filaments of ADP-actin than in monomers (Oda, Iwasa et al. 2009, Fujii, Iwane et al. 2010, von der Ecken, Müller et al. 2014, Galkin, Orlova et al. 2015) but the mechanism of hydrolysis was uncertain without structures of ATP- and ADP-P_i_-actin. Conformational changes in our filament structures bring Q137 and H161 closer to the γ-phosphate and must also reposition the two water molecules. Two configurations of the phosphate chain fit equally well into our EM map of AMPPNP-actin (Figs. 7C, D), so we propose two possible organizations of the waters and phosphate chain that could stimulate hydrolysis. Both models have Water 1 hydrogen bonded to both Q137 and H161 but differ in the orientation of Water 1 relative to the bond between γ-phosphorous atom and the bridging atom (N3B in AMPPNP or O3B in ATP). The model in Fig. 7D has Water 1 in line for attacking the γ-phosphate, while the alignment is imperfect in the model in Fig. 7C. Quantum mechanical MD simulations (McCullagh, Saunders et al. 2014, Sun, Sode et al. 2017) may be able to distinguish these alternatives. High-resolution crystal structures of ATP-actin in a filamentous conformation would also be very informative.

### How does phosphate release change dissociation of subunits from filament ends and the affinity of filaments for cofilin and other proteins?

Our structures of actin subunits buried in filaments revealed that the conformations of AMPPNP-, ADP-P_i_-and ADP-actin are remarkably similar. Nothing about the overall conformation provides clues about why phosphate dissociation changes the rates of subunit dissociation or the affinity for cofilin. However, the loss of hydrogen bonds and a divalent cation bond in the absence of the γ-phosphate might explain why ADP-actin filaments are more flexible than ATP or ADP-P_i_-filaments (Isambert, Venier et al. 1995).

Greater flexibility within a subunit may reduce its affinity for the end of a filament. For example, Narita et al. (Narita, Oda et al. 2011) found that the conformation of pointed ends of muscle actin filaments differs from the core of the filament. However, more information about the structure of filament ends will be required to connect subunit flexibility to the mechanism of subunit dissociation.

Greater flexibility alone may account for the higher affinity cofilin for ADP-actin filaments than ADP-P_i_-actin (Cao, Goodarzi et al. 2006). Cofilin is unlikely to bind to the standard conformation of polymerized actin, given that cofilin-decorated filaments have a tighter helical twist (163°) than undecorated filaments (167°) (McGough, Pope et al. 1997, Galkin, Orlova et al. 2011). Furthermore, the association rate constant for cofilin binding filaments is <1% the expected value (Blanchoin and Pollard 1999), so Blanchoin proposed that only 1% of the subunits of ADP-actin filaments are in the high energy (163°) conformation required to bind cofilin. Reconstructions of undecorated filaments from electron micrographs established that the subunits have a range of twist angles and that cofilin binding stabilizes a minor, high energy state (Galkin, Orlova et al. 2001). Cofilin binding to subunits with highly twisted conformations is an example of the conformational selection theory for protein interactions (Boehr and Wright 2008). Since subunits with bound ATP or ADP-P_i_ have more hydrogen bonds between the two halves of the molecule and those filaments are stiffer, the energy barrier between the equilibrium state and the 163° state will be much higher and thus less populated.

### Conformational changes in the actin polymerization cycle

Most of the >130 actin crystal structures in RCSB PDB are in a closed conformation, even though these actins came from different species and bound different nucleotides, ions and proteins (Fig. S10). Two exceptions (PDBs: 3ub5 and 1hlu) are actin bound to profilin in an open conformation (Chik, Lindberg et al. 1996, Porta and Borgstahl 2012). Actin subunits in filaments are similar in all three nucleotide states in a flattened conformation different from the closed and open conformations of monomers (Fig. S10). By combining structures of monomers and filaments with different bound nucleotides, we have the snapshots required to reconstruct conformational changes in the whole cycle of assembly and disassembly (Fig. S11).

## Materials and Methods

### Actin Purification and Polymerization

Muscle acetone powder was made using the flash-frozen chicken muscle from a local Trader Joe’s grocery store (MacLean-Fletcher and Pollard 1980). Actin was purified using one cycle of polymerization and depolymerization followed by gel filtration through Sephacryl S-300 and stored in Ca-G-Buffer (2 mM Tris-HCl, pH 8.0; mM ATP; 0.1 mM CaCl_2_; 1 mM NaN3; 0.5 mM DTT). Ca-ATP-actin was converted to Mg-AMPPNP-actin (Courtemanche and Pollard 2013), Mg-ADP-P_i_-actin and Mg-ADP-actin before being polymerized into filaments (Pollard 1986). We purchased ATP (A2383), AMPPNP (A2647), ADP (A2754) and hexokinase (H6380) from Sigma-Aldrich (St. Louis, MO), AG 1-X4 resin (1431345) from Bio-Rad (Hercules, CA) and glucose (167454) from Thermo Fisher Scientific (Waltham, MA).

### Sample Vitrification and Image Acquisition

The holy carbon C-flat 2/1 grids (Protochips company in Morrisville, NC) were used for all the cryo-EM samples. The grids were glow-discharged for 6 seconds in a Solarus plasma cleaner (Gatan company in Pleasanton, CA) in Ar (75%)/O2 (25%) at 25 W. To vitrify the samples, 3 μL of actin filament solution polymerized from 4 μM actin monomers was applied onto the carbon side of the grid in Mark III Vitrobot (FEI company in Hillsboro, OR) at 20°C at 95% of humidity. After incubating on the grid for 15 seconds, extra solution was blotted off using standard Vitrobot paper (grade 595; Ted Pella in Redding, CA) for 4.5 seconds at offset 1. Grid pre-screening was performed on an F20 microscope operated at 200 kV and equipped with a K2 Summit camera (FEI company). The three datasets were collected on a Titan Krios microscope equipped with an XFEG at 300 kV, a nanoprobe and a Gatan image filter (slit width: 20 eV). Image stacks were recorded on a K2 Summit camera in super-resolution mode, controlled by SerialEM (Mastronarde 2003). For AMPPNP-actin and ADP-P_i_-actin, a Volta phase plate was used at a defocus of −0.5 μm (Danev, Tegunov et al. 2017) and the dose rate was set to ~3.8 counts/pixel/second. For ADP-actin, image stacks were recorded at a defocus value between −1 and −2.5 μm, without the phase plate, and the dose rate was set to ~8.0 counts/pixel/second. For all three datasets, each image was fractionated into 32 frames (0.25 seconds/frame), and the physical pixel size was 1.045 Å.

### Image Processing

Dose-fractioned image stacks were dose-averaged, magnification-corrected, motion-corrected, and summed with MotionCor2 (Zheng, Palovcak et al. 2017) using 9 × 9 patches. The frames in the first second of image recording (with large drifts), and those in the last second for ADP-actin (with high radiation damage) were discarded. CTF parameters were estimated with Gctf (Zhang 2016) using the unweighted sums. Filaments were manually boxed out with sxhelixboxer.py in SPARX (Hohn, Tang et al. 2007). The filaments coordinates were exported from SPARX and imported into RELION2 (He and Scheres 2017) for further analysis. Filaments were windowed into square segments using a box size of 328 × 328, and an interparticle distance along the long axis of 26 pixels. First, we worked on a subset (~20,000 particles) of the whole dataset for each sample (~120,000 particles). Following 2D classifications, we reconstructed 3D maps using the known helical parameters (rise: 27.3 Å; twist: −166.5°) and a simulated model, in which each actin subunit is depicted as a ball. The maps from small datasets were filtered to 10 Å before being used as reference models for reconstructions using the whole datasets. After post-processing, all three maps went beyond 3.8 Å resolution at this point. Later we performed 2D and 3D classifications to remove bad particles (~10%). The well-resolved classes from each 3D classification were similar. Particles in good classes of 2D and 3D classifications were pooled together for final reconstructions. A soft-edged 3D mask with a radius of 45% of the box size was created for post-processing. The B-factors for map sharpening were first determined by RELION2 itself. Later, we used a slightly bigger (less negative) B-factor than the one determined by RELION2. The B-factors for AMPPNP-actin, ADP-P_i_-actin, and ADP-actin were −75, −75 and −100 (Å^2^), respectively. The Fourier Shell Correlation 0.143 criteria (FSC_0.143_) were used for resolution estimation. Layer-line images were calculated from map projections with SPARX (project and periodogram commands) (Hohn, Tang et al. 2007). Local resolutions were calculated with ResMap (Kucukelbir, Sigworth et al. 2014). We used both micrograph CTFs and particle CTFs for reconstructions, and the estimated resolutions were almost the same. All the image processing was carried out on Yale High Performance Computing servers.

### Model Building and Refinement

The atomic models were built with Coot (Emsley, Lohkamp et al. 2010). Most of residue side chains were built unambiguously. When there was an ambiguity, we referred the local conformations of the corresponding residue in the 1.5 Å resolution crystal structure of rabbit actin (PDB: 1j6z) (Otterbein, Graceffa et al. 2001). The primary sequence of chicken actin is the same as that of rabbit actin. Refinements were carried out for several rounds in reciprocal space with REFMAC (Brown, Long et al. 2015) and then in real space with PHENIX (Afonine, Poon et al. 2018). The models from REFMAC had slightly better fitting statistics, while the models from PHENIX had better geometry as analyzed with Coot. At last, we chose the structures from PHENIX as the final models.

### Structure Analysis and Presentation

The inter-domain rotation angles were calculated with the DynDom web server (Hayward and Lee 2002). The rise (translation) and twist (rotation) for the helices were calculated with Chimera (match showMatrix command) using two inter-strand adjacent subunits in models, which are pixel-size independent. RMSDs were also calculated with Chimera (rmsd command) (Huang, Couch et al. 1996). The structural figures were generated with MolScript/Raster3D (Merritt and Bacon 1997) (Fig. S1A), PyMOL (Schrodinger 2015) (Fig. S6), and Chimera (Huang, Couch et al. 1996) (rest of structural figures)

## Acknowledgements

Research in this publication was supported by National Institute of General Medical Sciences of the National Institutes of Health under awards number R01GM026132 and R01GM026338. The content is solely the responsibility of the authors and does not necessarily represent the official views of the National Institutes of Health. The authors thank Dr. Shenping Wu, Dr. Marc Llaguno and Dr. Xinran Liu for managing the microscopes, Yale High Performance Computing support team for maintaining the clusters, Moon Chatterjee and Shoshana Zhang for commenting on the manuscript and members of our lab for helpful discussions.

## Supplemental materials

**Figure S1.**
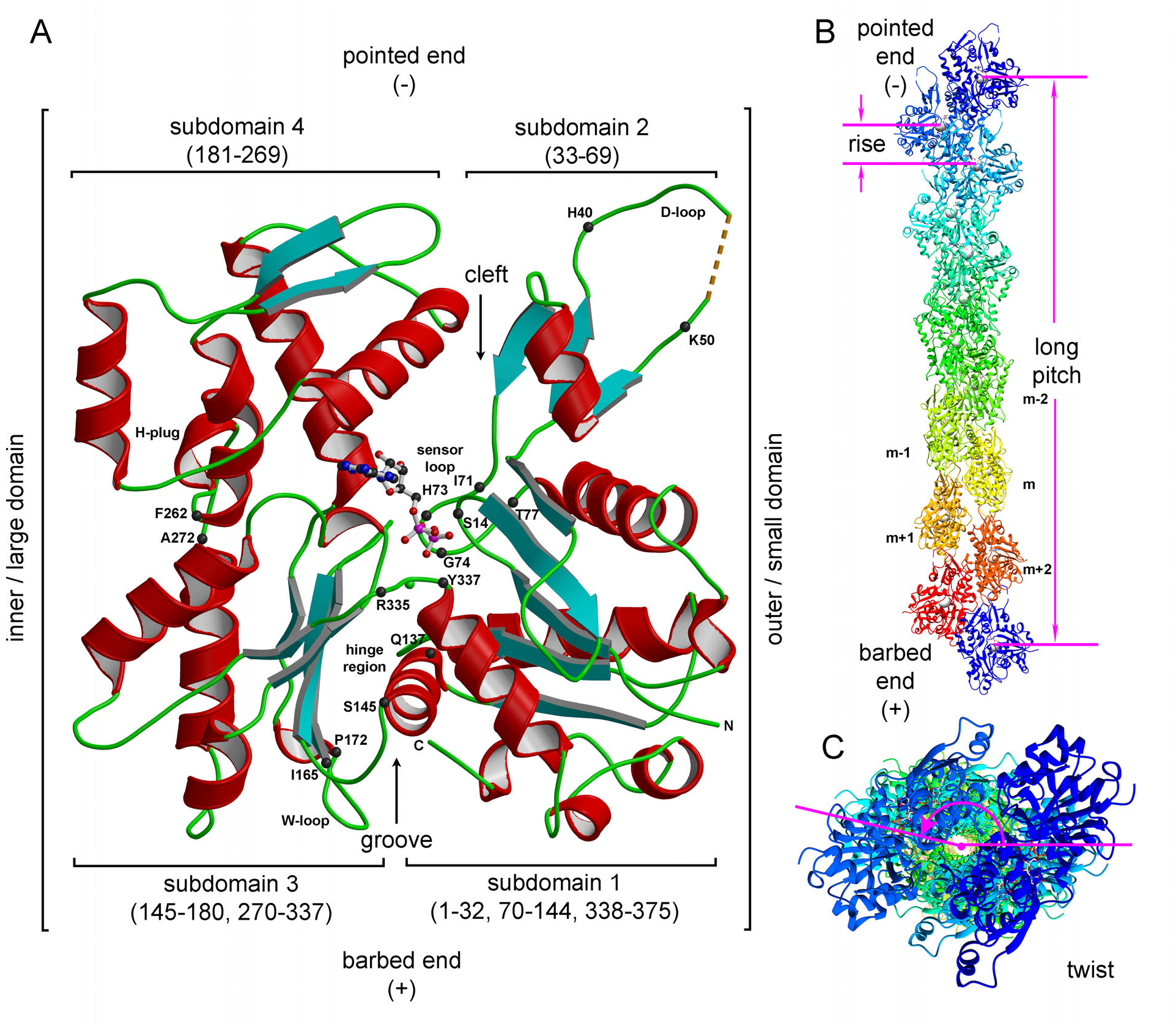
Structural elements of actin subunits and helical assembly of actin filaments. (A) Structural elements on an ADP-actin subunit. The DNase I-binding loop (D-loop; residues 40-50), sensor loop (residues 71-77), WH2-domain binding loop (W-loop; residues 165-172), hydrophobic plug (H-plug; residues 262-272) and hinge region (hinge helix: residues 137-145; hinge loop: residues 335-337) are highlighted. (B) Helical assembly of ADP-actin filament with 14 subunits (side view). The grey ball in each subunit indicates the center of mass of the subunit, which is very close to β-phosphate of the ADP molecule. The rise per subunit along the short pitch helix is 27.52 Å and half a turn along the long-pitch helix is ~360 Å. The barbed end is facing downward. (C) Top view (from pointed end to barbed end) of ADP-actin filament. The twist for our ADP-actin filament is −166.63 ° (minus sign: left-handed helix).

**Figure S2.**
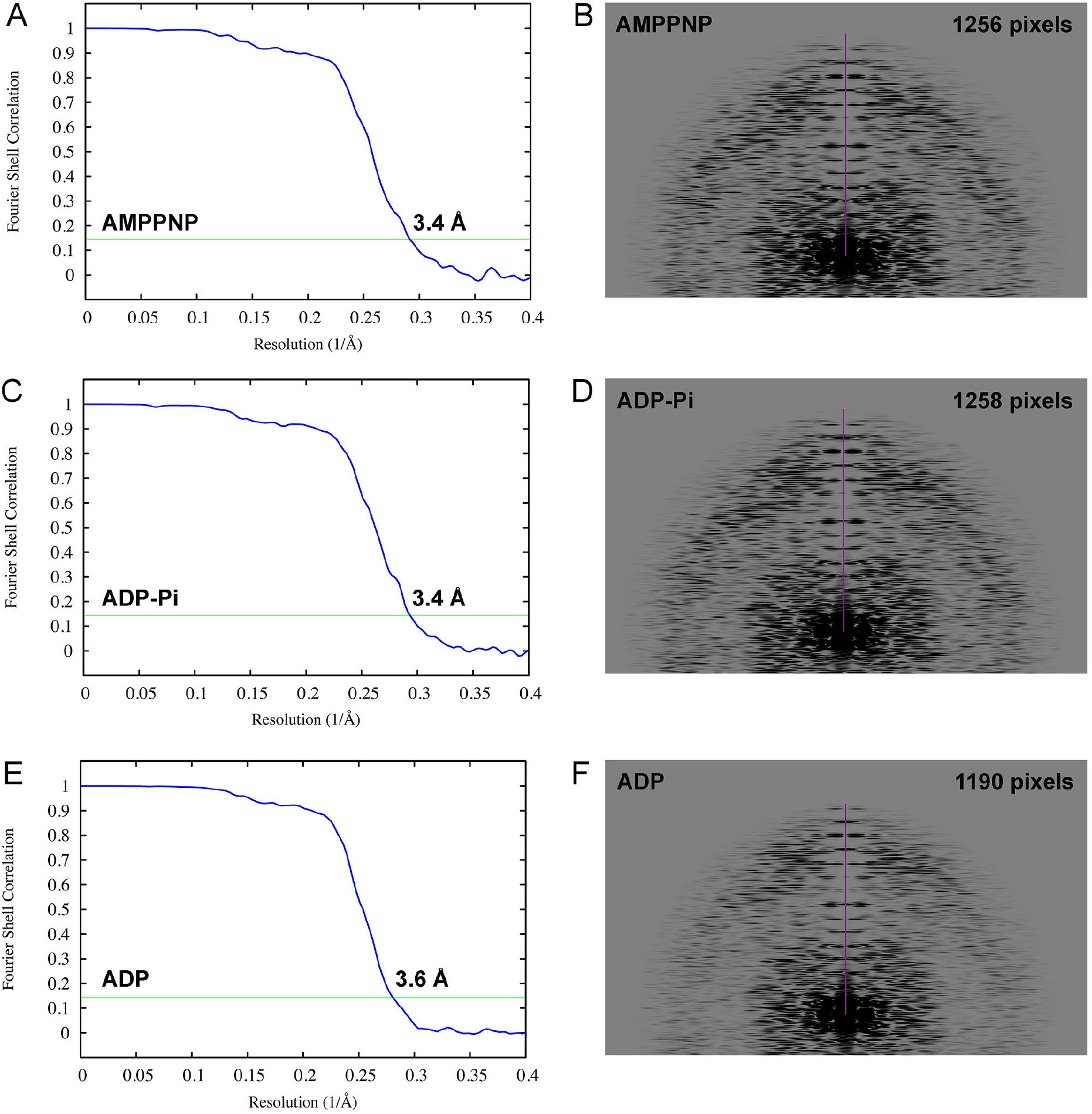
Global map resolution estimation. Resolution estimation of (A) AMPPNP-, (C) ADP-P_i_-, and (E) ADP-actin reconstructions using Fourier Shell Correlation (FSC; blue curves) with 0.143 criteria (green horizontal lines). The resolutions are indicated in unit of angstroms. Resolution estimation of (B) AMPPNP-, (D) ADP-P_i_-, and (F) ADP-actin reconstructions using layer-line images calculated from back projected images. The purple vertical line are the heights from the origin to the highest visible layer line in units of pixels. The layer-line images are 4096 × 4096 pixels, and the pixel size of back projected images is 1.045 Å. The resolutions are 3.4 Å for AMPPNP-actin filaments, 3.4 Å for ADP-P_i_-actin filaments and 3.6 Å ADP-actin filaments using the formula: resolution = (pixel size) × (layer-line image size) / (layer-line height).

**Figure S3.**
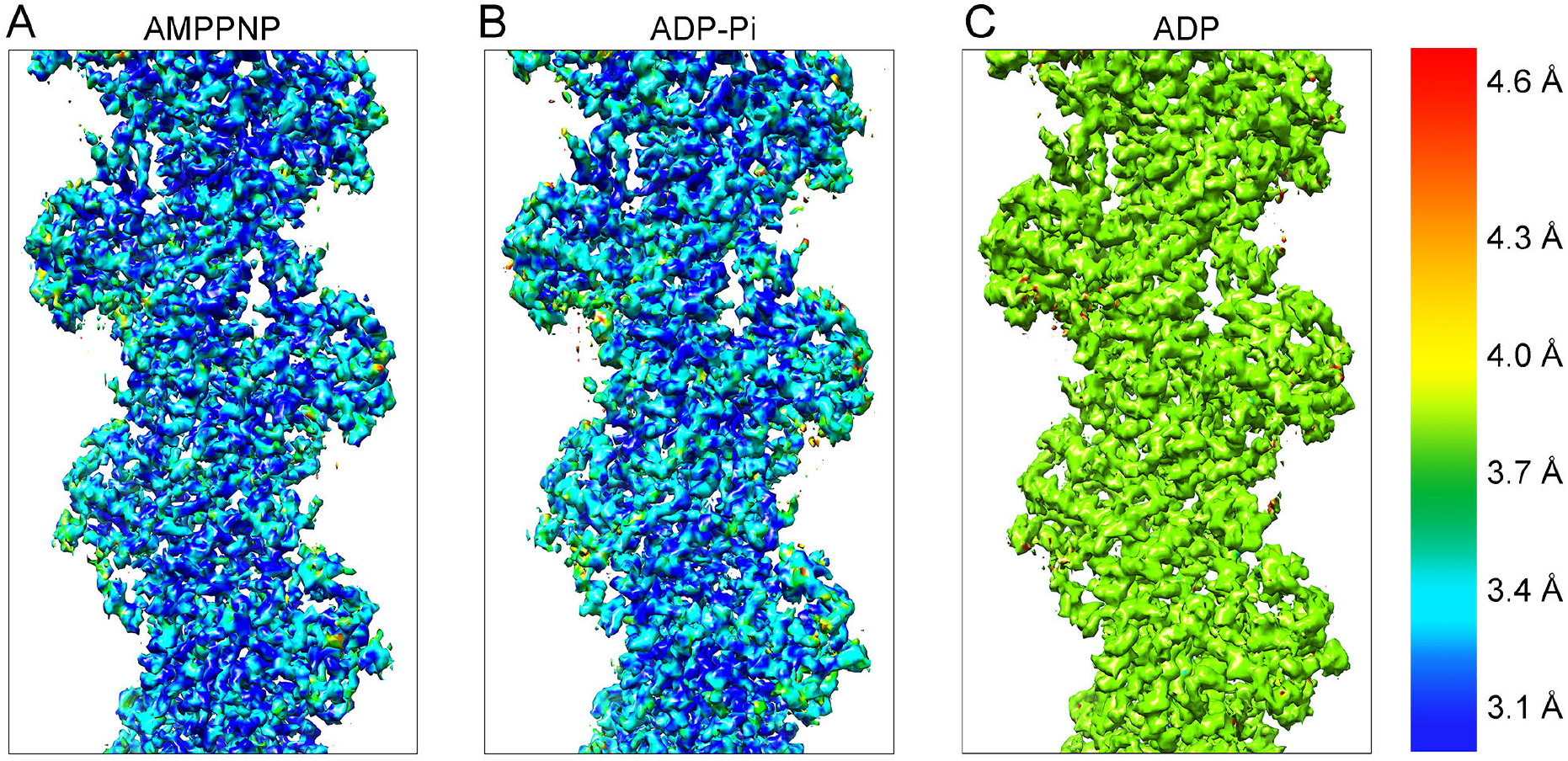
Local resolution estimated by ResMap of the three actin filament reconstructions mapped onto the cryo-EM densities. (A) AMPPNP-actin, (B) ADP-P_i_-actin, and (C) ADP-actin. The color-coded scale is on the right side.

**Figure S4.**
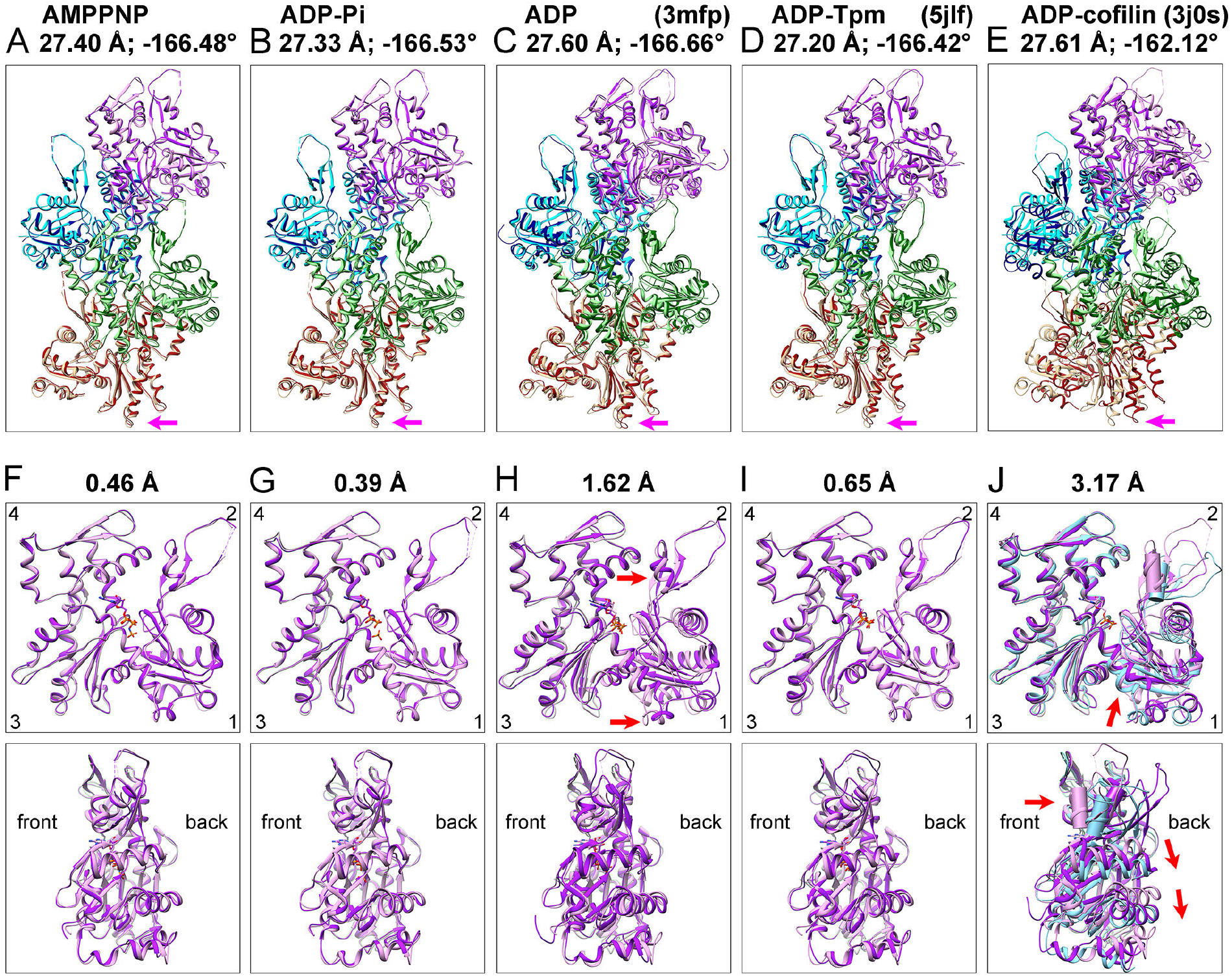
Ribbon diagrams comparing our ADP-actin filament model with six actin filament structures. The ADP-actin filament has a rise (subunit translation) of 27.52 Å and twist (subunit rotation) of −166.63° (minus sign: left-handed helix). (A-D) Superimpositions of four subunits from our ADP-actin filament model with the comparison filaments. The upper subunit of the ADP-actin filament was aligned with the upper subunit of each structure: (A) our AMPPNP-actin; (B) our ADP-Pi-actin; (C) ADP-actin (PDB: 3mfp); (D) tropomyosin-decorated ADP-actin (PDB: 5jlf); and (E) ADP-actin saturated with cofilin (PDB: 3j0s). The subunits in our ADP-actin structure are light colors (plum, cyan, light green and tan) and subunits in other structures are dark colors (purple, blue, dark green and dark red). The pink arrows point to a loop in subdomain 3 of the 4^th^ (tan) subunit to show differences in subunit translation and rotation. (F-J) Superimpositions of the first subunit of our ADP-actin model with the first subunit in (A-E) after alignment of the Cα atoms in subdomains 3 and 4 (residues 145-337). The number at the top of each panel is the RMSD of Cα atoms of residues 5-370 relative to our ADP-actin structure, excluding the flexible regions (residues 1-4 and 45-49) in the N-terminus and D-loop. (J) Includes the crystal structure of an actin monomer (PDB: 2a42) in light blue. The only helix in subdomain 2 (residues 55-61) is rendered as a cylinder to compare inter-domain rotation angles. The red arrows point to structural differences (upper panel) and directions of motion (lower panel).

**Figure S5.**
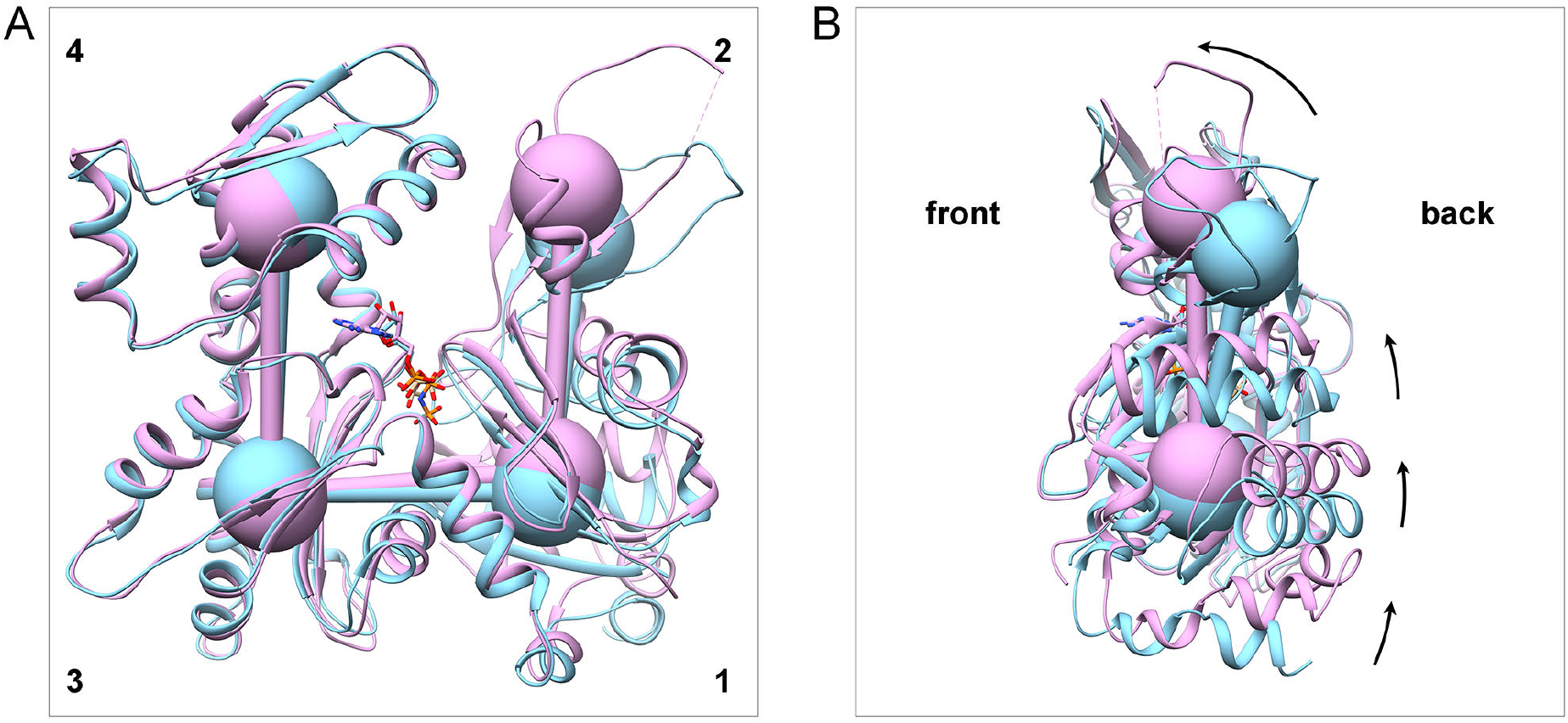
Inter-domain rotation and translation. (A) Front view and (B) side view of actin molecules. One actin subunit in our AMPPNP-actin filaments (plum) and one rabbit ATP-actin monomer (light blue) (PDB: 2a42) are superimposed after aligning subdomains 3 and 4. The balls indicate the centers of mass of subdomains. The primary sequences of these two molecules are the same.

**Fig. S6.**
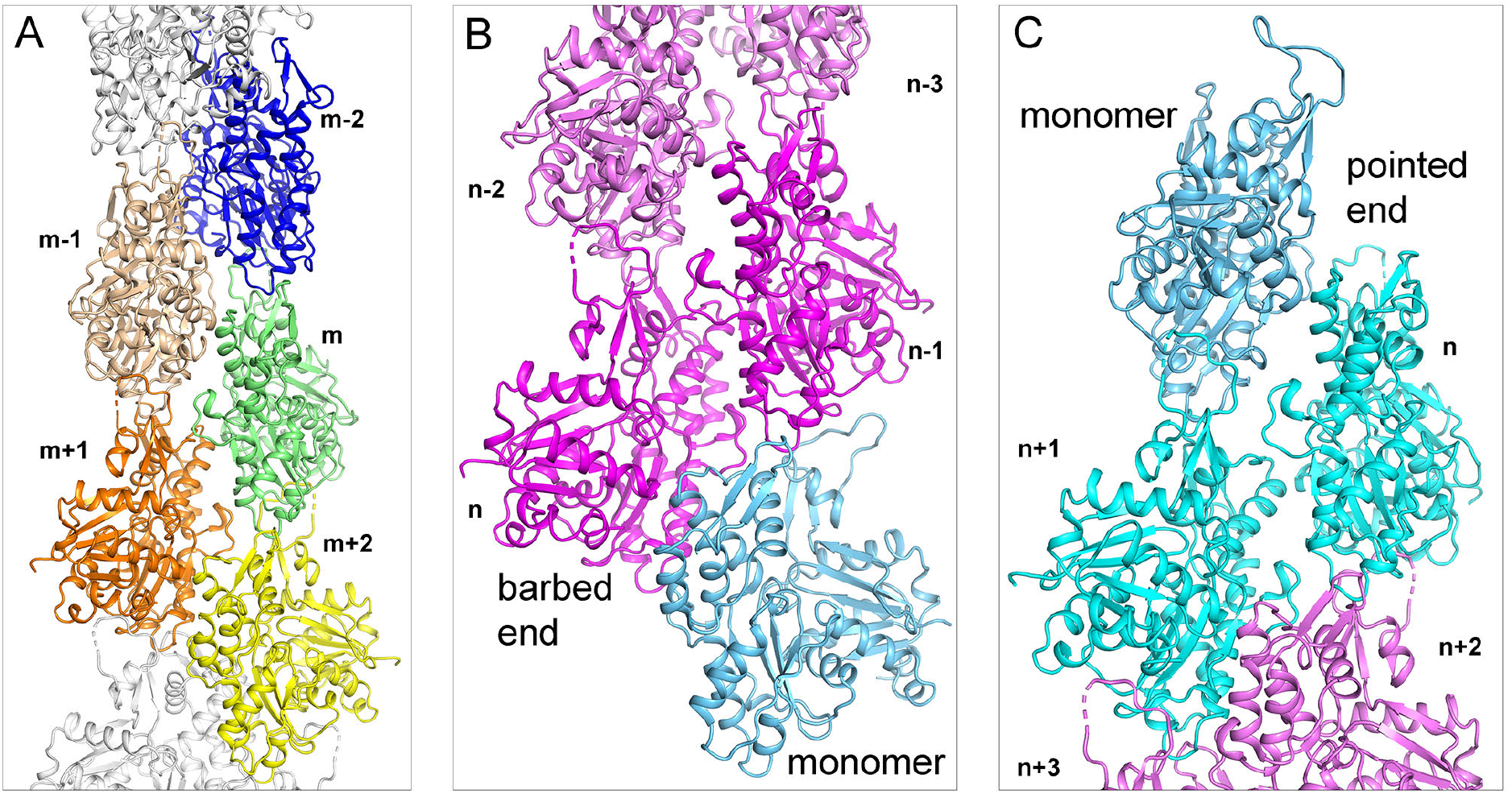
Ribbon diagrams showing subunit interactions in filaments and possible conformations of subunits at the two ends. (A) Overview of filament showing contacts along the long-pitch helix (*m-2* to *m* to *m+2*) and interstrand contacts with two subunits along the short-pitch helix (*m-1* to *m* to *m+1*). (B) At the barbed end subunits *n* and *n-1* (magenta) likely have conformations similar to subunits in the middle of the filament (plum), because longitudinal contacts (between n and n-2, and between n-1 and n-3) flatten these two subunits. The flattened conformation of these barbed end subunits is favorable for interactions with the pointed end of an incoming monomer (light blue). (C) At the pointed end, subunits *n* and *n+1* (cyan) are likely in a conformation similar to monomers with disordered D-loops, because longitudinal contacts (between *n* and *n+2* and between *n+1* and *n+3*) cannot flatten these two subunits. These conformations are unfavorable for interactions with an incoming action monomer, which is also in the unflattened conformation.

**Figure S7.**
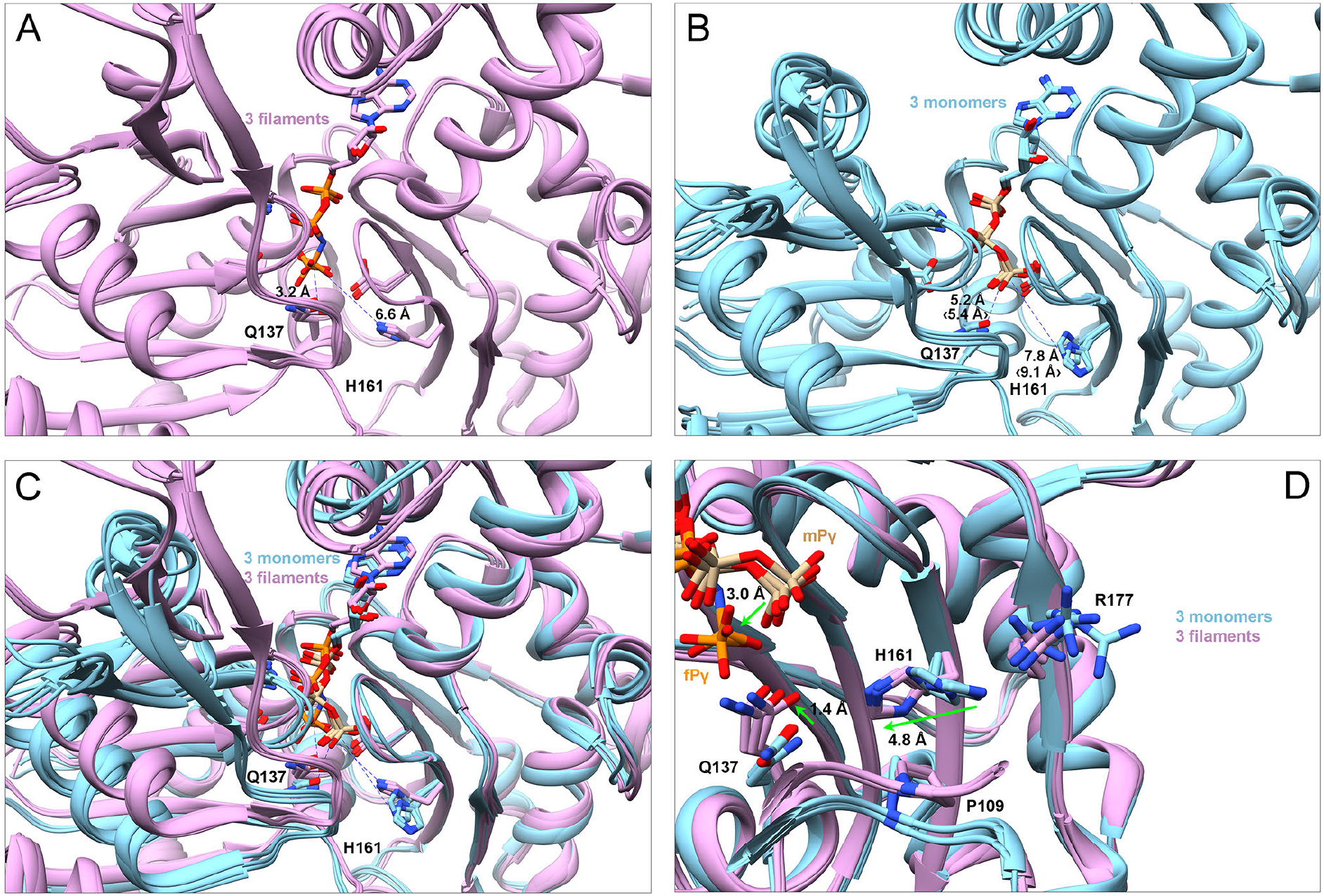
Ribbon diagrams with stick figures of the nucleotides and crucial side chains in the active site during the ATPase cycle of polymerized actin. All the molecules are aligned using subdomain 3 (residues 145-180 and 270-337) and all the filament subunits in (A) are in the same orientation as they in Fig. 5. Phosphorus atoms in filaments are colored in orange; those in monomers in tan. (A) Superimposition of one subunit from AMPPNP-, ADP-P_i_- and ADP-actin filaments. In the AMPPNP actin filament, the γ-phosphorous is 3.2 Å from the side chain OE1 atom of Q137 and 6.6 Å from the side chain NE2 atom of H161. (B) Superimposition of ATP-actin monomers from rabbit (with bound Ca^2+^ and DNase I, PDB: 2a42), *Dictyostelium* (with bound Li+ and human gelsolin subdomain 1; PDB: 1nmd) and budding yeast (with bound Mg^2+^ and human gelsolin subdomain 1; PDB: 1yag). We flipped the imidazole ring in 1nmd, based on an analysis of water network. In 1nmd, the γ-phosphorous atom is 5.2 Å from the side chain OE1 atom of Q137 and 7.8 Å from the side chain NE2 atom of H161. In other monomeric structures, these two distances are about 5.4 Å and 9.1 Å. (C) Superimposition of all the actin molecules in (A) colored plum and (B) colored light blue. (D) Side view (in an orientation similar to Fig. 3B) showing the relative movements of residues in the catalytic center during polymerization (green arrows). The γ-phosphate group is bent downward in filaments (fP_γ_).

**Figure S8.**
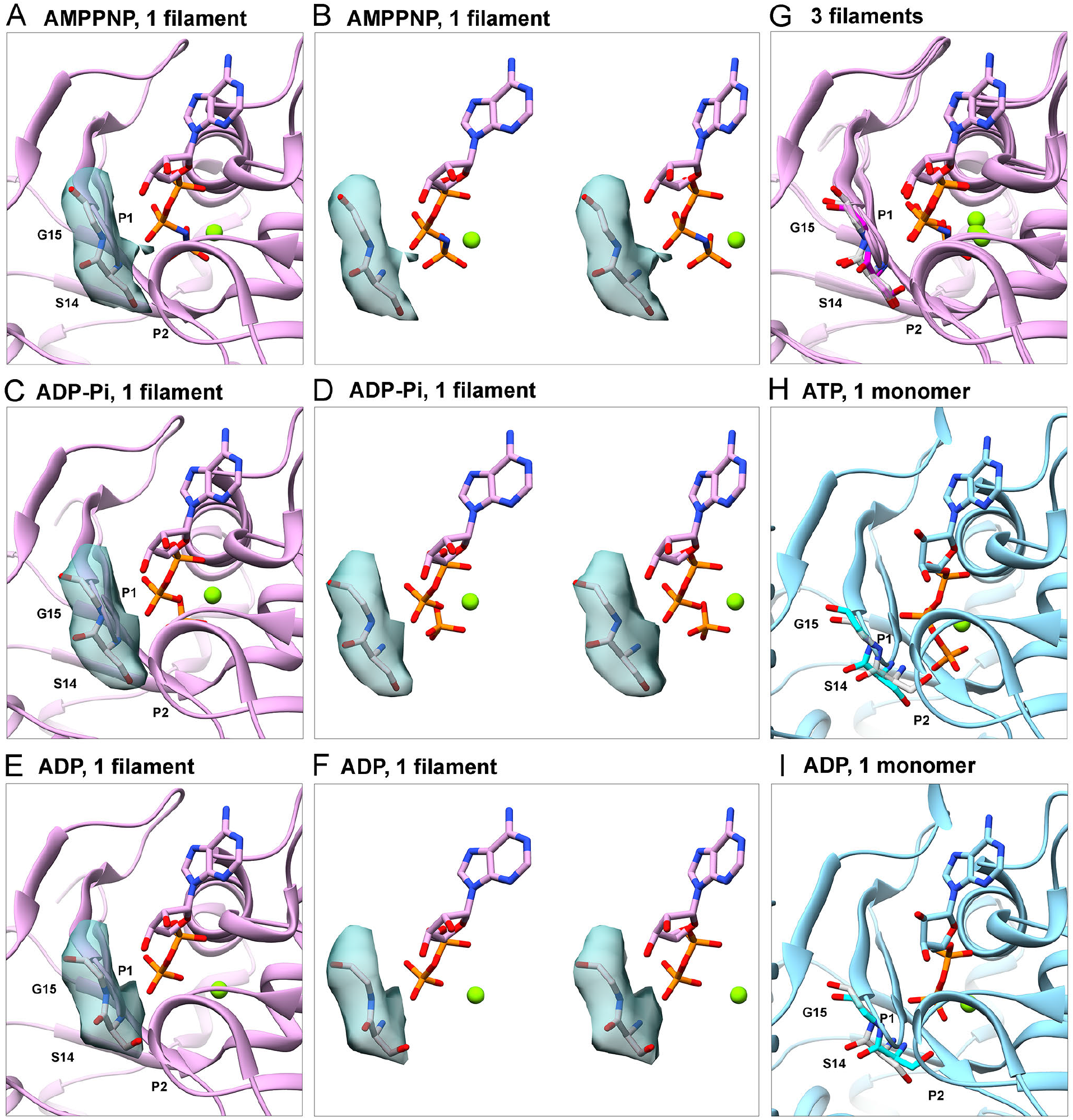
S14 is a direct sensor of nucleotide states. (A), (C) and (E) Ribbon diagram of subunits in filaments of AMPPNP-, ADP-P_i_- and ADP-actin. Residues S14 and G15 are shown as stick figures with their EM surface maps. (B), (D) and (F) Wall-eye stereo stick figures of S14, G15, the nucleotide and Mg^2+^ (green ball) and the EM surface maps of the residues. (G) Superimposition of AMPPNP-, ADP-P_i_- and ADP-actin filament subunits. S14 and G15 are rendered as semi-transparent grey stick figures in AMPPNP-and ADP-P_i_-actin, and as a magenta stick figure in ADP-actin. (H) Ribbon diagram of an actin monomer with bound ATP and Mg^2+^ (PDB: 1nm1). S14 and G15 are rendered as cyan (PDB: 1nm1) and semi-transparent grey (PDB: 3a5l, bound with ADP and Mg^2+^) stick figures. (I) Ribbon diagram of an actin monomer bound with ADP and Mg^2+^ (PDB: 3a5l). S14 and G15 are rendered as cyan (PDB: 3a5l) and semi-transparent grey (PDB: 1nm1, bound with ATP and Mg^2+^) stick figures. All the structures are aligned using subdomain 3 and displayed in the same orientation. Local EM maps are contoured as in Fig. 5.

**Figure S9.**
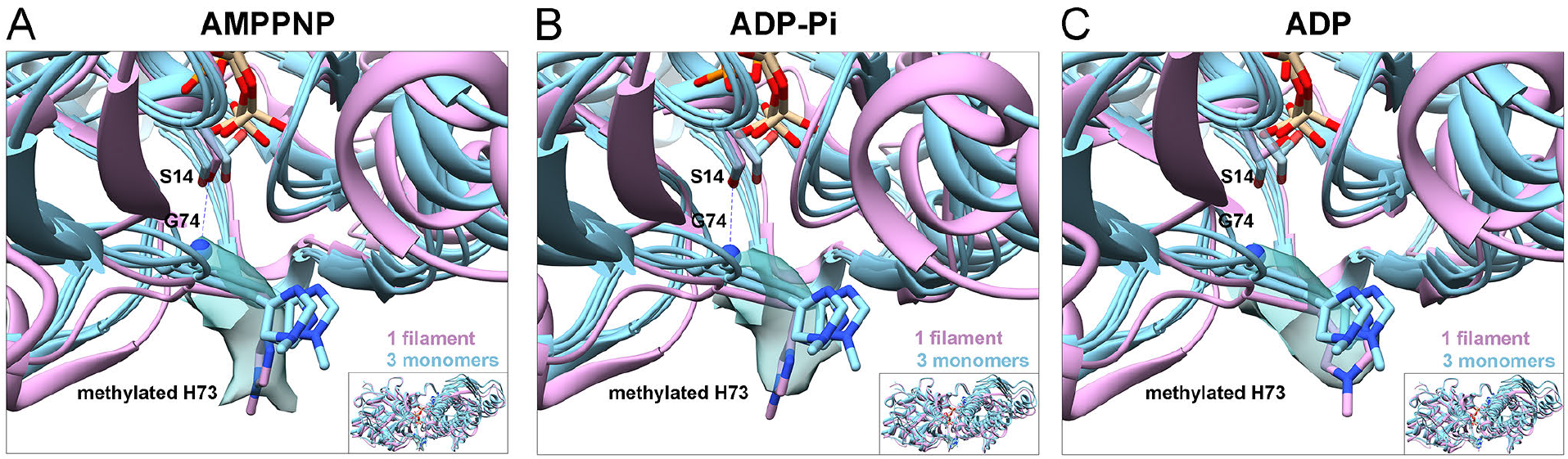
In filaments, the side chains of methylated H73 and S14 respond to nucleotide states synergistically through the hydrogen bond between S14 and G74. (A-C) The map densities of methylated H73 in AMPPNP-, ADP-P_i_-, and ADP-actin filaments shown as surfaces and stick figures of the models in plum. Three actin monomers (light blue) from rabbit [PDB: 2a42; Ca-ATP], *Dictyostelium* [PDB: 3a5l; Mg-ADP] and budding yeast [PDB: 1yag; Mg-ATP]) are superimposed to the filament model (plum) of AMPPNP-, ADP-P_i_-, or ADP-actin by aligning subdomain 1. The G74 backbone nitrogen atom is rendered as a blue ball. The side chains of S14 in monomers are shown as semi-transparent light blue sticks. Phosphorus atoms in filaments are colored in orange; those in monomers in tan. The maps of H73 in the structures of AMPPNP-actin and ADP-P_i_-actin are contoured at 0.043 e/Å^3^, and the map of H73 in the structure of ADP-actin is contoured at 0.055 e/Å^3^. The spikes around H73 in AMPPNP-actin filaments could be due to noise or multiple conformations.

**Fig. S10.**
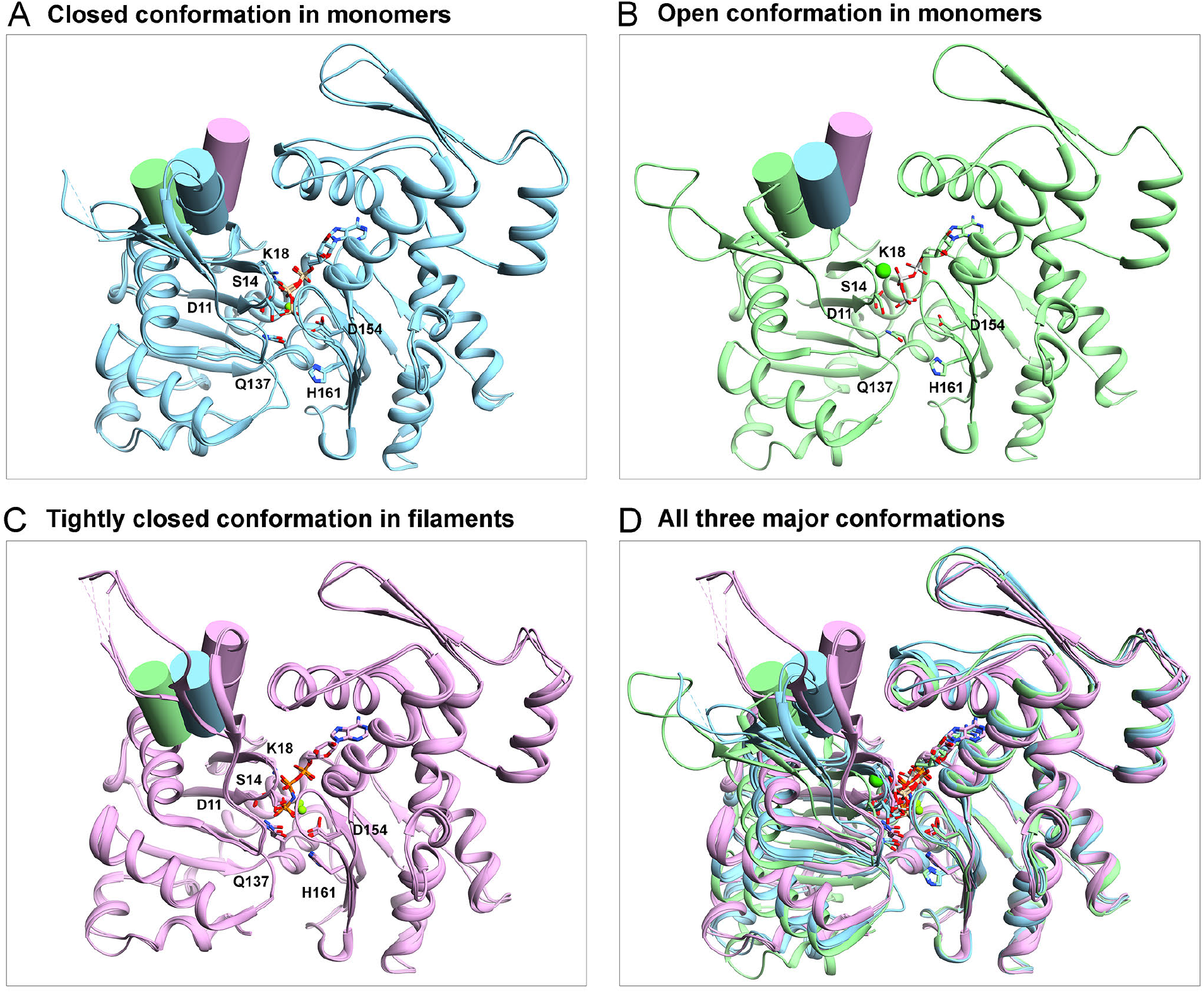
Three major conformations of actin molecules. Structures are aligned using subdomain 3 (residues 145-180 and 270-337). (A) Superimposition of ATP-*(Dictyostelium* actin complexed with Mg^2+^ ion and human gelsolin segment 1; PDB: 1nm1) and ADP-*(Dictyostelium* actin complexed with Mg^2+^ ion and human gelsolin segment 1; PDB: 3a5l) actin monomers showing the closed conformation. (B) Ribbon diagram of ATP-actin monomer (bovine β-actin complexed with Ca^2+^ ion and bovine profilin; PDB 3ub5) showing the open conformation. (C) Superimposition of our AMPPNP-, ADP-P_i_, and ADP-actin in filaments showing the tightly closed conformation. (D) Superimposition of all six structures from (A), (B) and (C). The only helix (residues 55-61) in subdomain 2 is rendered as light blue (in closed conformation), light green (in open conformation) or plum (in tightly closed conformation) cylinder.

**Figure S11.**
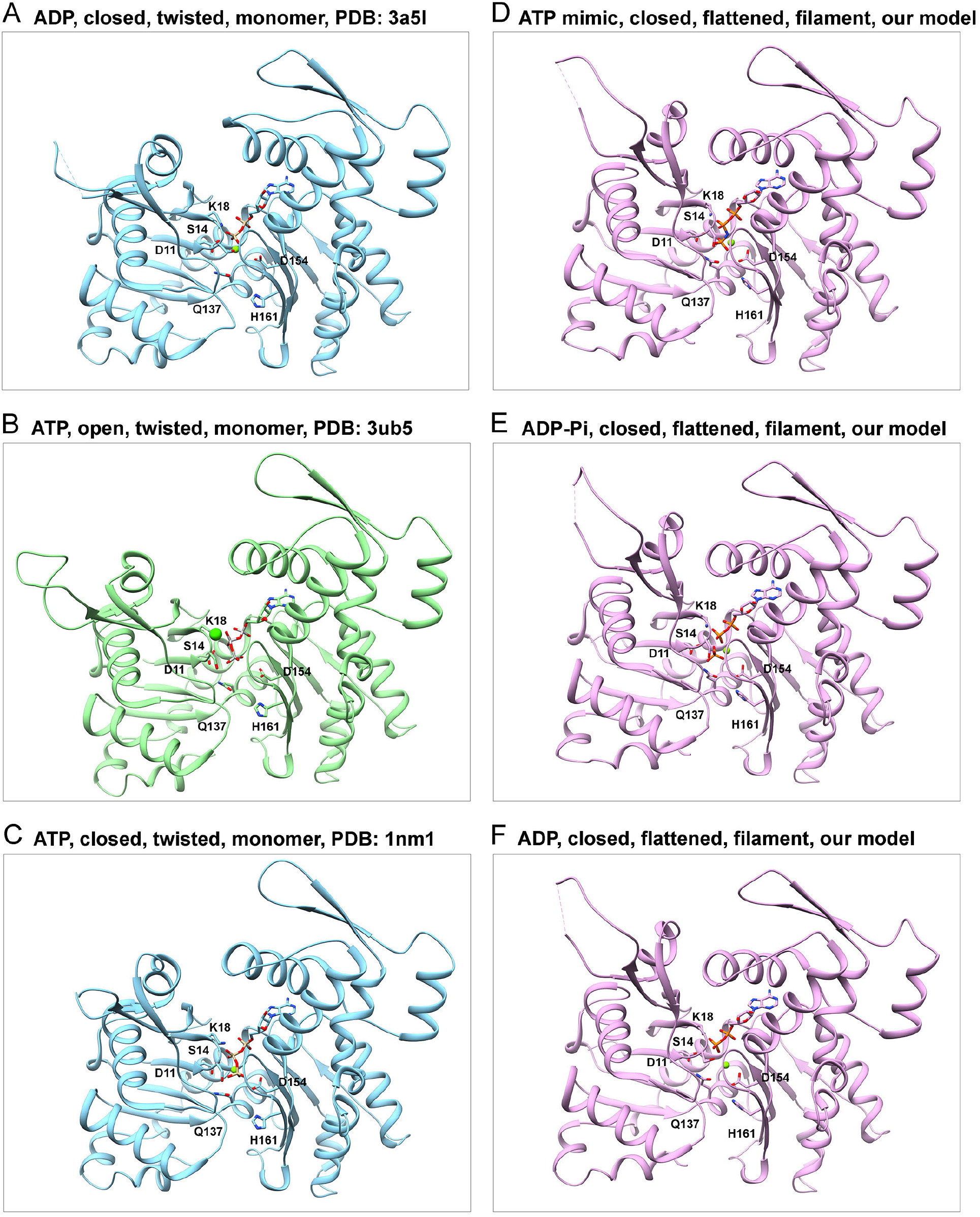
The cycle of conformational changes in the actin molecule during the cycle of ATP binding, polymerization, ATP hydrolysis and phosphate release illustrated by the six structures in Fig. S10. (A) Closed conformation of the ADP-actin monomer (Otterbein, Graceffa et al. 2001). In the absence of the γ-phosphate (P_γ_), the direct sensor of nucleotide state in the P1 loop, S14, is hydrogen-bonded to the backbone of G158 (in the P2 loop). The distance between OG atom of S14 and N atom of G158 is 3.1 Å, appropriate for a hydrogen bond. Compared with ATP actin monomers, the P1 loop collapses slightly toward the hinge helix (Figs. S1 and S8I). (B) Open conformation of the ATP-actin monomer bound to profilin (Chik, Lindberg et al. 1996, Porta and Borgstahl 2012). The nucleotide-binding cleft is open, increasing the rate of ADP dissociation by 14 fold (Vinson, De La Cruz et al. 1998). ATP binds rapidly to nucleotide-free actin (De La Cruz and Pollard 1995). (C) Closed conformation of the ATP-actin monomer. With bound ATP the side chain of S14 rotates by ~120° (a different rotamer) to form a hydrogen bond with the backbone of G74 in the center of the sensor loop (residues 71-77) (their distance: 2.6 Å) (Kabsch, Mannherz et al. 1990, Otterbein, Graceffa et al. 2001, Vorobiev, Strokopytov et al. 2003). The P1 loop moves upward away from the hinge helix by ~1.1 Å. (D) Flattened conformation of a subunit in the AMPPNP-actin filament brought about by (i) inter-domain rotation (Figs. S5, 3A) (Oda, Iwasa et al. 2009), (ii) inter-subdomain rotation (domain flattening) (Fig. 3 B and C) (Fujii, Iwane et al. 2010), (iii) local conformational changes in W-loop, D-loop and C-terminal tail (Fig. 5), and (iv) rearrangements in the active site that move the side chain of Q137, the imidazole ring of H161 and water 1 to attack the γ-phosphate (Figs. 6 and 7). (E) Flattened conformation of a subunit in the ADP-P_i_-actin filament. The structure is virtually identical to the AMPPNP-actin filament, except that the hydrolyzed P_γ_ moves a small distance from P_β_ (Fig. 5A, B). (F) Flattened conformation of a subunit in the ADP-actin filament. P_i_ is released slowly through the back channel (Wriggers and Schulten 1999). P1 loop moves slightly towards the hinge helix to occupy some space vacated by P_i_ release, and the side chain of S14 establishes new contact with the backbone of G158 (Figs. S8 and S9). The imidazole ring of H73 moves slightly closer to subdomain 3 after P_i_ release (Fig. S9).

